# Regenerative base editing enables deep lineage recording

**DOI:** 10.64898/2026.02.06.703857

**Authors:** Duncan Chadly, Ron Hadas, Leslie Klock, Jiahe Yue, Felix Horns, Amjad Askary, Alejandro Granados, Remco Bouckaert, Carlos Lois, Long Cai, Michael B. Elowitz

## Abstract

Reconstructing the lineage histories of individual cells can reveal the dynamics of developmental and disease processes. In engineered recording systems, cells stochastically edit synthetic barcode sequences as they proliferate, creating distinct, heritable edit patterns that can be used to reconstruct the lineage trees relating individual cells in a manner analogous to phylogenetic reconstruction. However, recording depth is often limited by the kinetics of the editing process: the rate of editing declines exponentially over time for an array of independently editable targets, leading to most edits occurring in early generations. Here, we introduce the hypercascade, a regenerative molecular recording system that takes advantage of the predictability of A-to-G base editing to progressively create new target sites over time. The hypercascade packs 4 editable target sites in every 20 bp of sequence, enabling high density information storage. More importantly, the hypercascade’s regenerative logic leads to an approximately constant rate of mutation accumulation over time. This in turn facilitates reconstruction of deep lineage relationships. We demonstrate this by reconstructing trees spanning 23 days of editing and approximately 17 generations after a single polyclonal engineering step. Finally, simulations show that the hypercascade has the potential to record chromatin state transition dynamics across multiple genomic loci in parallel. The hypercascade thus provides a flexible and broadly useful tool for molecular recording.

## Introduction

Multicellular organisms typically develop from a single cell through repeated cycles of division and differentiation. Taken together, the set of division events form a lineage tree that can reveal the impact of lineal history on cell fate potential^1,2^, provide insight into diseases involving the generation of pathological cell division patterns^3^ Therefore, identifying the lineage relationships between cells has been a fundamental goal across both developmental biology and medicine.

Despite their centrality, lineage trees have been difficult to map in most organisms and contexts. Individuals can exhibit substantial variability in their underlying cellular lineage processes^4^. In classic work, researchers mapped the deterministic cellular lineage tree of *C. elegans* through direct observation.^5^ However, other organisms typically present greater challenges: they are optically opaque, preventing direct time lapse imaging; they can be much larger, comprising many thousands or even millions of cells in a tissue or context of interest; the lineages that form tissues are variable, limiting the ability to integrate observations across multiple individuals^4^.

To address these challenges, several groups have engineered synthetic lineage recording systems, such as MEMOIR^6,7^, GESTALT^3,8,9^, CARLIN^10^, LINNAEUS^11^, SMALT^12^, homing CRISPR barcoding^13,14^, and others^15–17^. These systems use different methods, including recombinases^7^, CRISPR nucleases^3,6,8–11,13–17^, and CRISPR base editors^12^ to dynamically edit (mutate) specific, heritable target sequences as cells proliferate. Each cell lineage thereby accumulates a unique edit pattern. Edits can subsequently be read out in bulk or, ideally, in individual cells, either by sequencing^3,8–16^ or microscopic imaging^6,7,17^. Finally, computational algorithms allow reconstruction of lineage relationships from measured endpoint edit patterns, in a manner loosely analogous to phylogenetic reconstruction. Most recently, barcode labeling by incorporating temporally ordered sequences using Prime Editing technology has been introduced as a new paradigm for phylogenetic recording.^17–20^

In this work, we sought to address two remaining key difficulties with current lineage recording methodologies that constrain the depth (number of cell generations) of recording. First, in most recording systems, edit rates are proportional to the number of unedited target sites. This causes the fraction of unedited sites to decay exponentially over time, rather than at a constant rate^21^. Frontloading mutations in this way consumes the majority of memory capacity early, with little ability to record information later in the process. Second, the total amount of memory (editable sites) available in most recording systems can be small (e.g. tens of editable target sites^3,6–10,16,19^). This further limits the potential duration of recording and resolution of resulting trees.

To address these issues, we designed a new generative recording system termed the hypercascade. This system takes advantage of A-to-G base editors^22^, which can make specific, single base pair mutations with high fidelity and allow dense packing of target sites. The defining feature of the hypercascade is that editing of existing target sites generates new editable target sites in an approximately 1:1 ratio. Edits can therefore accumulate over time without diminishing the number of remaining editable target sites for an extended period. In simulations, this generative property linearizes the rate of editing over time and improves the accuracy of lineage reconstruction. This strategy also enables packing of target sites at a high density of approximately one target site every 5 base pairs of sequence, allowing a large amount of memory to be integrated into single cells for recording.

Experimental analysis in mouse and human cells showed that the generative editing property works as designed. One-shot engineering of the system into human induced pluripotent stem cells (hiPSCs) enabled lineage reconstruction of over 3000 cells, identifying clonal phenotypes. Finally, based on experimental measurements of the effects of chromatin modification on edit rate, we show that the hypercascade has the potential to enable recording of chromatin state transitions over time. Future work will aim to develop multiplexed single-cell methods to fully take advantage of the strengths of hypercascade recording.

## Results

### The hypercascade design allows sustained recording through generative editing

In contrast to some editing mechanisms, CRISPR A-to-G base editors (ABEs) are distinguished by their ability to generate precise and predictable single base mutations at defined target sites (**Figure 1A**). Further, tandem arrays of base editor target sites allow compact genetic encoding of multiple bits of recorded information. Edits can accumulate over multiple cell generations and are stably inherited by daughter cells (**Figure 1B**). Consequently, readout of edits by sequencing can in principle be used to reconstruct cell lineage relationships (**Figure 1B**).

**Figure 1:**
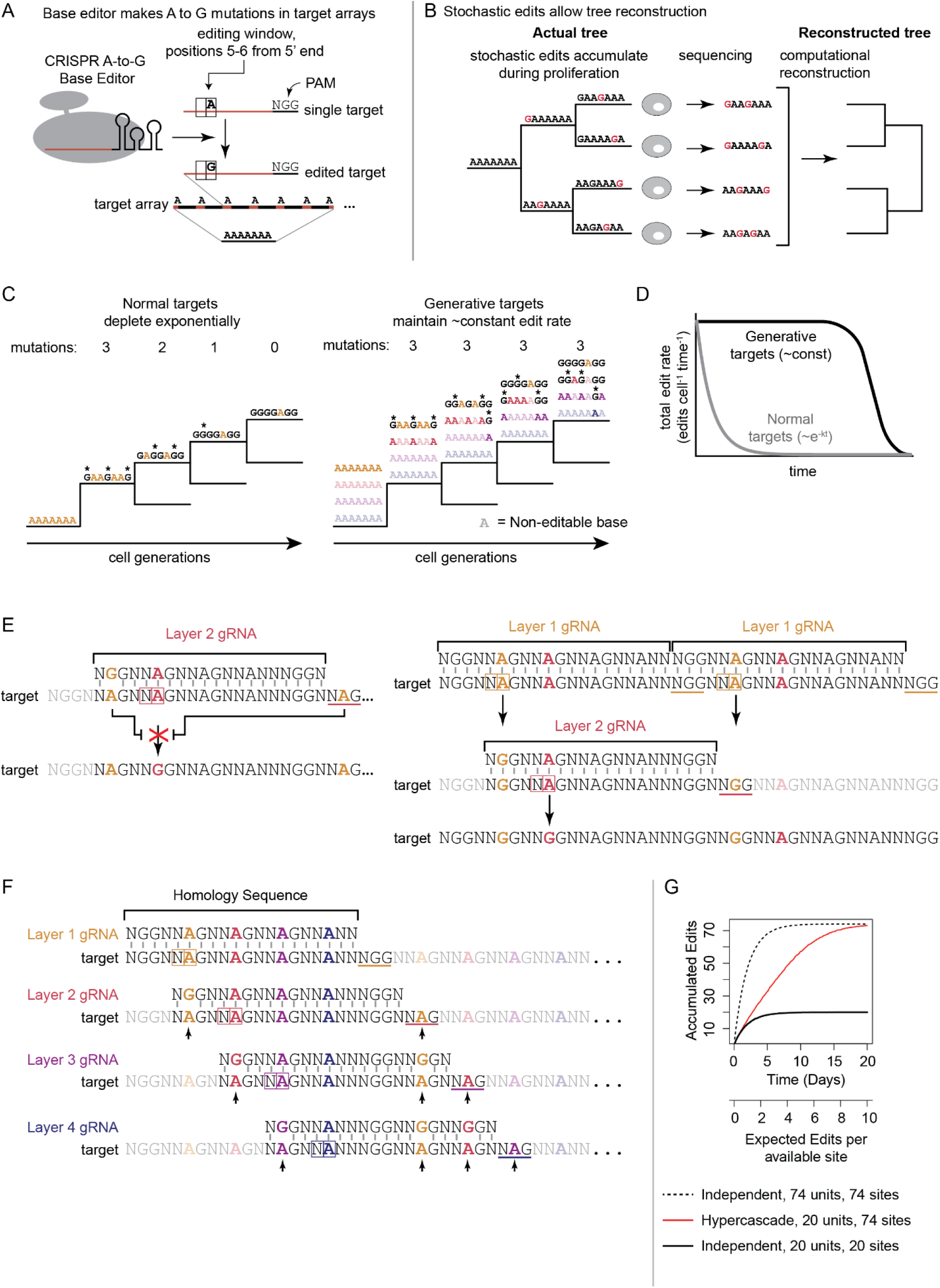
The hypercascade system linearizes edit rate over time and densely packs mutable target sites. **(A)** The CRISPR A-to-G base editor makes predictable mutations at defined target sites. **(B)** Reading out heritable mutations enables reconstruction of cellular lineage relationships. **(C)** Arrays of independently edited targets are exponentially lost over time; in contrast, a system that generates new targets over the course of editing would maintain constant edit rate over an extended time scale. **(D)** Generative targets are predicted to maintain approximately constant edit rate until the generative mechanism fails (schematic). **(E)** New targets can be generated by repairing protospacer and PAM mismatches through A-to-G mutations. **(F)** This concept can extend to multiple unlocking layers of densely packed target sites. This sequence consists of a tandem repeating 20-mer which is acted on by four unique guide RNAs. Mismatches that are repaired to generate new targets indicated with arrows. **(G)** Simulations of editing in this scheme reveal linear edit accumulation relative to arrays of independent targets.

Even if it densely encodes multiple bits, a simple tandem array of base editable target sites faces a fundamental limit in its recording capacity: if each individual site has an independent and constant probability of editing per unit time, as has been observed^21^, then the number of available unedited sites decays exponentially over time. Early generations receive many edits while later generations receive few or none (**Figure 1C, D**).

One way to circumvent this problem is to generate new target sites as old ones are consumed by editing. As long as new sites can be generated, this scheme could keep the total number of unedited sites, and therefore the edit rate, approximately constant over time. For example, imagine a set of target sites organized into 4 logical “layers” (**Figure 1C**, right panel) such that only layer 1 target sites (upper row) are initially accessible to editing. Further, assume that editing of layer 1 sites generates editable layer 2 sites (second row). Similarly, editing of layer 2 sites could generate layer 3 sites (third row), and layer 3 edits could generate layer 4 sites (fourth row). In such a scheme, editing would consume and generate sites at roughly the same rate, keeping the total edit rate constant over time until nearly all potential edits have occurred (**Figure 1D**).

To implement this scheme and provide high density memory encoding, we designed a base editable target sequence, termed the hypercascade, that exploits unique properties of ABEs. The ABE has two requirements for efficient function. One is a 20 bp homology sequence (encoded by the gRNA it complexes). The other is a 3 bp protospacer adjacent motif (PAM), which is NGG for the most commonly used Cas9 homolog derived from *S. pyogenes*.^22^ When these two requirements are met, an A in the fifth or sixth position of the target sequence is mutated to a G (**Figure 1A**).^22^ The predictability of this edit outcome makes it possible for editing of available target sites to repair engineered mismatches that otherwise prevent editing by distinct gRNAs at other target sites (**Figure 1E**).

Taking advantage of this edit-repair principle, we designed a cascading system that packs four target sites, representing four logical “layers,” into one tandemly repeatable 20bp sequence. These target sites can be edited using four corresponding gRNAs. However, only the layer 1 target site starts with a perfect match to its cognate gRNA and a functional PAM. Each of the others is initially prevented from editing by at least two mismatches in its protospacer sequence and PAM (**Figure 1F**).

The final feature of this design is intended to enforce an ordered sequence of edits (**Figure 1F**). Editing of a layer two site only becomes possible after two surrounding layer one edits. Similarly, editing of a layer 3 site requires previous editing of two surrounding layer 2 sites, and a previously edited layer 1 site. Finally, editing of each layer 4 site requires previous editing of two surrounding layer 3 sites, as well as previous editing of the layer 1 and 2 sites necessary for those layer 3 edits. We created an animated video to explain and visualize the operation of the system at youtu.be/GRVMbn-dElc. Overall, this design allows dense packing of editable bases while requiring only four distinct gRNAs for operation. Specification for this scheme leaves 11 base pairs unconstrained in the 20bp repeating element, yielding a total of approximately 4 million possible designs. In analogy to the way the term hypersphere generalizes the notion of a sphere, we term this structure, which comprises a 2-dimensional array of linked cascades, a hypercascade (**Figure 1E, F**).

To test whether the hypercascade could address the exponential memory decay problem, we simulated a stochastic edit process using the Gillespie algorithm^23^, and compared a hypercascade of 20 repeating units (74 target sites total in a 403bp sequence) to two simpler independent control editing schemes. The first contained 20 edit sites, one per 20bp unit, comprising a similar overall DNA length (403bp), which is convenient for standard Illumina sequencing readouts. The second control scheme contained 74 non-interacting target sites, all of which were initially available for editing, which could be encoded in a longer stretch of 1.48kb of DNA. Thus one control scheme maintained the same DNA length while the other contained the same number of target sites.

With either of the control schemes, the number of subunits available for editing decreased exponentially over time, as expected (**Figure 1G**). By contrast, in the simulated hypercascade, edits accumulated at a nearly constant rate. For a given edit rate, the hypercascade extends the duration and linearity of recording relative to independent recorders with the same total DNA length, which saturate earlier regardless of how many total target sites they contain. Thus, simulations indicate that the hypercascade should successfully extend editing, maintaining an approximate linear accumulation of edits over a longer period of time compared to independent arrays of similar length.

### Simulations show that hypercascades reconstruct lineages more accurately than independent arrays

The hypercascade is intended to function as a lineage recorder. Therefore it is critical to test not only whether edits occur linearly over time but also whether typical edit patterns can allow accurate lineage reconstruction. To address this question, we simulated stochastic editing of either the 74 site hypercascade or one of two independent control target arrays in a population of cells undergoing repeated rounds of division (**Figure 2A**). In all systems we scanned a range of edit rates, array copy numbers, and lineage tree depths. After simulating trees in the forward direction, endpoint barcode states were used to reconstruct cell lineage relationships using the Unweighted Pair Group Method with Arithmetic mean (UPGMA)^24^. Finally, reconstructed trees were compared to ground truth trees using the normalized Robinson-Foulds distance^25^, where 0 represents a perfect topological match between the ground truth and reconstructed trees, and 1 represents two maximally different trees (**Figure 2A**).

**Figure 2:**
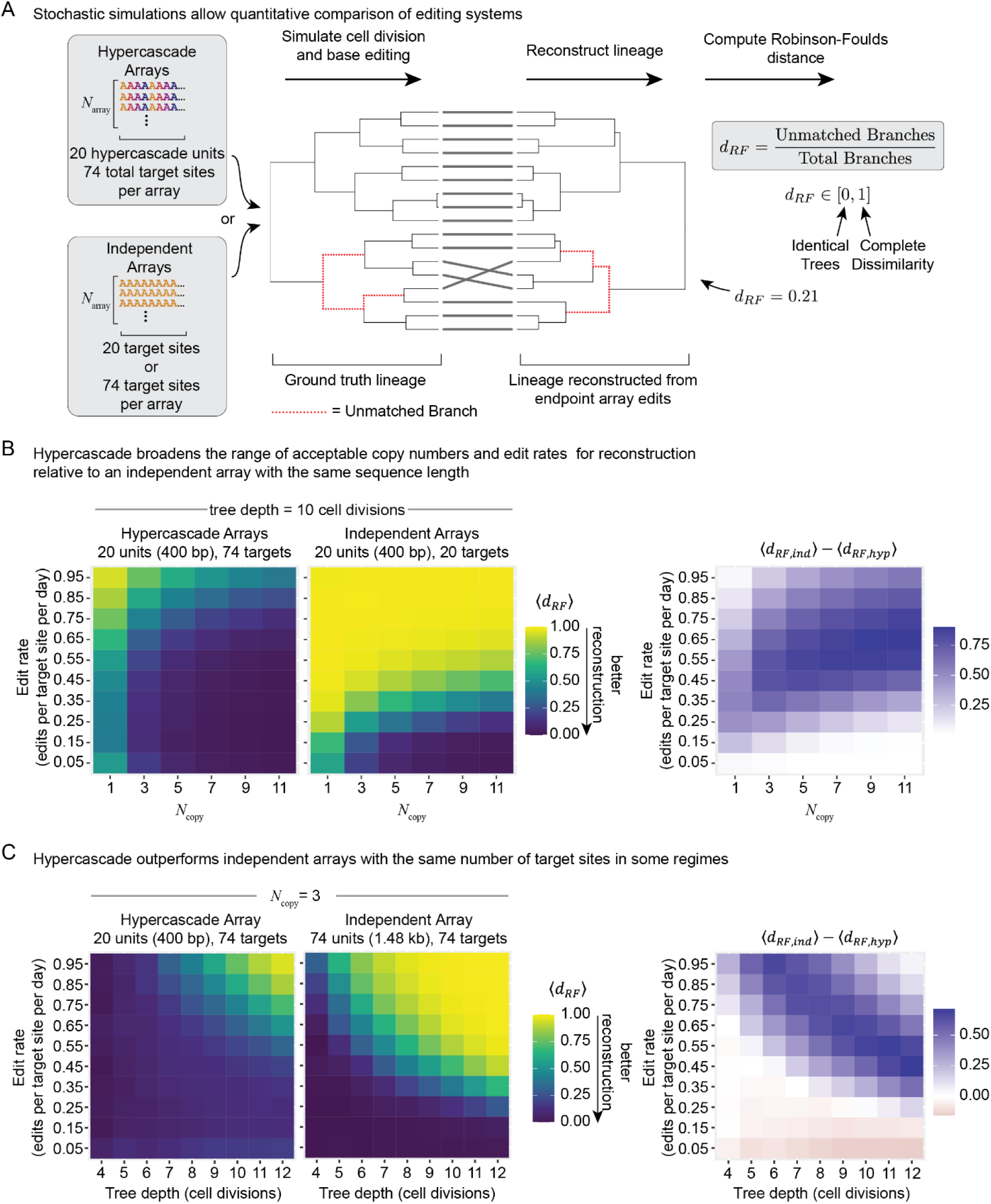
Hypercascades allow lineage recording and reconstruction across a broader range of target copy numbers and edit rates compared to arrays of independent sites. **(A)** Stochastic simulations of target editing and cell division together enable comparison of lineage reconstruction accuracy between independent and hypercascading systems. **(B)** Holding tree depth constant, the hypercascade system outperforms an independent array of targets with comparable sequence length over a variety of edit rates and copy numbers. **(C)** Hypercascading arrays also outperform independent arrays with the same total number of targets across a range of parameters, especially when edit rates are high or trees are large. For very slow edit rates, when both systems are far from saturation, independent arrays with comparable total targets moderately outperform the hypercascade due to entropic favorability.

We first compared the hypercascade to an independent array of equal DNA length. We fixed the mean tree depth at 10 cell divisions and varied both edit rate and array copy number (**Figure 2B**). At low edit rates, the hypercascade performed comparably to a 20-unit array. This is expected, as the first layer of the hypercascade is equivalent to 20 independent units. At higher edit rates, two benefits of the hypercascade became clear. First, high reconstruction accuracy was attainable for fewer integrations of the hypercascade compared to the 20-unit array (**Figure 2B**). Second, the hypercascade was relatively insensitive to edit rate. This feature is enabled by the generation of new target sites as old sites are depleted, linearizing the loss of editable targets over time (**Figure 1D**). Similar results were obtained using a simple model of Cas9 editing that produces multiple terminal edit outcomes at each of the 20 target sites (**Supplemental Figure 1**). Thus, for the same total DNA length and edit rate, the hypercascade is either similar to or outperforms the independent target arrays.

We next compared the hypercascade to an independent editable system with the same total number of target sites (**Figure 2C**). Because fully independent targets are uncorrelated, they have the potential to contain more information (higher entropy) than hypercascade targets, where some barcode states are never realized due to the ordered nature of editing. We fixed the target copy number to 3 and varied both edit rate and mean tree depth in both simulated systems. At very low edit rates, the independent array modestly outperformed the hypercascade. However, in regimes of higher edit rate or tree depth, where independent arrays saturate, the hypercascade dominated due its extended editing dynamics (**Figure 2C**). We expect that the hypercascade would also outperform larger independent arrays for deeper trees, even when edit rates are low, due to earlier saturation in the independent case.

### Different hypercascade sequences operate orthogonally with distinct kinetics

To assess the functionality of the hypercascade in cells, we constructed three hypercascade sequence arrays, all based on the same design principles, but differing at unconstrained sequence positions (**Figure 3A**). Each sequence contained 19 tandem repeats of its 20bp unit, plus an additional partial unit containing a PAM site, for a total length of ∼400bp, which is short enough to fit on a single sequencing read. To choose bases at the unconstrained positions, we used a machine learning model of Cas9 edit rates to select bases predicted to allow high edit rates in all four layers^26^ (**Figure 3A, Methods**).

**Figure 3:**
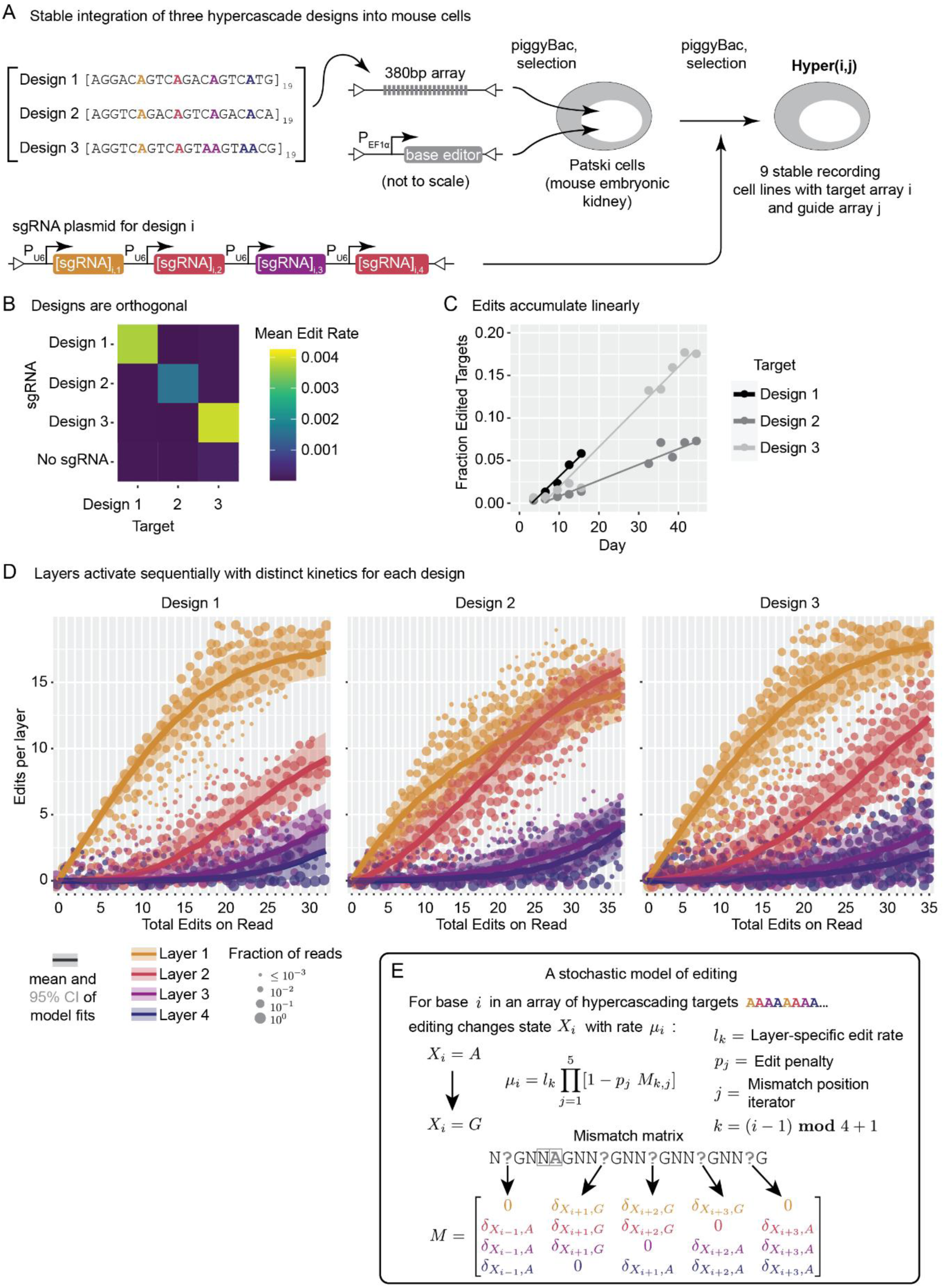
The hypercascade exhibits sequential editing in living cells. **(A)** Three different implementations of the hypercascade were stably integrated into a mouse embryonic kidney line with either on- or off-target gRNAs. **(B)** Editing occurs only in the presence of on-target gRNAs. **(C)** Edits accumulate linearly over a 44-day period. **(D)** Layers of targets sequentially activate and are well fit by a stochastic model of editing **(E)**.

We integrated each of the barcode arrays together with ABE in a mouse embryonic kidney fibroblast cell line^27^ via piggyBac transposition. We selected stable polyclonal lines, each containing ABE and one or more copies of the integrated hypercascade. We then initiated editing by transfecting piggyBac transposons containing gRNA expression constructs.

To test the orthogonality of the gRNA-target site pairs, we examined all pairwise combinations of gRNAs and target designs. We allowed editing to continue for up to 44 days after gRNA transfection, periodically harvesting cells, and sequencing their hypercascades. In these experiments, editing occurred only for the expected gRNA-hypercascade match, as expected (**Figure 3B**). Further, the fraction of edited sites increased linearly over time for all three target sequence designs (**Figure 3C**). These results indicate that hypercascade sequences provide a rich design space that could allow for multiple orthogonal recording channels in the same cell.

Previous studies with wild type Cas9 have shown that the nuclease edits targets with a small number of mismatches, although at a reduced rate, especially at targets with functional PAM sites.^46,47^ This mismatch editing efficiency has been found to depend on both the sequence of the protospacer and the distance of the mismatches from the PAM site, with higher tolerance of mismatches at sites further from the PAM.^46,47^ We therefore sought to quantify the existence and rate of potential out-of-order editing. Edit rates were highest for gRNAs with perfect homology with their target. Consistent with previous findings, the editing rate typically decreased with more mismatches and was especially suppressed when the PAM site was broken (**Supplemental Figure 4**).

Finally we asked whether edits accumulated sequentially through the designed mechanism (**Figure 1E**). We compiled reads from all time points, grouped them by the total edits per read and by their layer identity, and plotted the distribution of edits for each layer (**Figure 3D**). Each site was parameterized by a layer-specific edit rate, with multiplicative edit rate penalties for each possible mismatch, disfavoring premature edits. These plots revealed an ordered structure, with edits generally occurring in the expected sequence. Further, the patterns of accumulated edits were qualitatively well-fit by a stochastic model of editing incorporating all edit rate parameters, including the low rates of editing at mismatched target sites (**Figure 3D**, solid lines and **Figure 3E**; **Supplemental Figure 4**). Taken together, these results indicate that the hypercascade successfully generates new targets during editing, as designed, leading to the intended linear and sequential accumulation of edits.

### One-shot transfection of the hypercascade editing system reveals broad clonal features

A practical challenge for any recording system is integrating it into cells. In most systems, integration requires laboriously engineering cells to express multiple components, screening monoclones, and potentially repeating the whole process until cell lines with the desired characteristics are generated. This provokes the question of whether a complete hypercascade system could be integrated in a single step through piggyBac transposition. If so, resulting clones could then be selected in a single antibiotic selection step prior to downstream experiments being immediately performed on the resulting polyclonal cell line. This approach could enable recording in systems that are traditionally difficult to engineer or prone to silencing, including primary cells and hiPSCs.

We investigated this approach using hypercascade target sequence 1. We simultaneously transfected all components into hiPSCs, selecting with antibiotic for 1 day, then continuing culture of the cells and collecting samples every passage for genomic DNA extraction once they recovered (**Figure 4A**). After an initial delay, edits accumulated over time (**Figure 4B**). This delay may reflect excess episomal plasmid arrays persisting from the initial transfection, slowly diluting out over time, and competing with genomically integrated target sites for editing and amplification. After the initial delay, edits accumulated through day 23 post transfection (**Figure 4B**). As in the previous experiment (**Figure 3D**), the distribution of edits into different layers was consistent with sequential activation of layers (**Figure 4C**). These results indicate that generative hypercascade editing can be achieved with a single genome engineering step.

**Figure 4:**
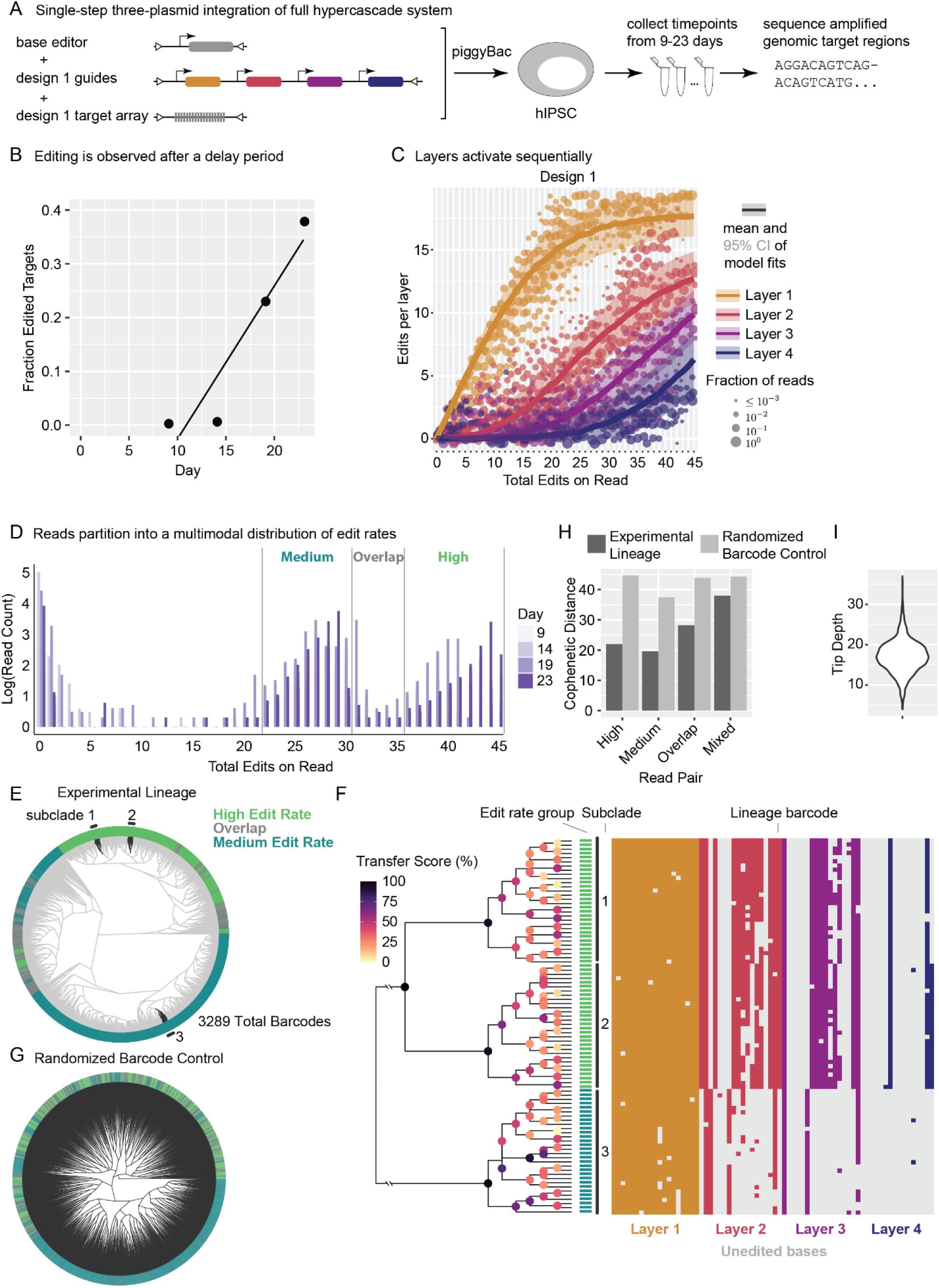
The hypercascade can be integrated in a single step and used for subsequent lineage recording. **(A)** All components of the system were simultaneously integrated into hiPSCs. **(B)** Edits accrued on integrated targets over a 23 day time course. **(C)** Layered targets sequentially activated. **(D)** Read counts binned by total edits present a multimodal distribution with peaks that shift right over time. **(E)** UPGMA-reconstructed lineage relationships between 3289 unique barcode sequences observed with at least 22 accumulated edits. **(F)** Uncertainty in the lineage was assessed for a subset of taxa using Bayesian phylogenetics (BEAST2). A subtree constructed from the taxa highlighted in **(E)** is visualized with transfer bootstrap scores annotated on each clade within the tree. **(G)** Randomization of barcode sequences destroys the experimentally observed tree structure. **(H)** The mean cophenetic distance between pairs of cells from distinct edit rate groups is lower than in the randomized barcode control, identifying edit rate as a clonal feature. **(I)** The average tip depth is estimated to be approximately 17 generations, consistent with experimental expectation.

Interestingly, the frequency of reads with different total numbers of edits naturally grouped into several distinct distributions (**Figure 4D**). These likely represent individual clones or groups of clones that received specific copy numbers and integration sites for the editing component plasmids, leading to different edit rates. Distributions shifted to the right over time, as would be expected in the context of genomic edit accumulation.

To reconstruct lineage relationships, we identified unique edit patterns within the data, leaving out targets with few edits that contain relatively little recorded information (**Figure 4E**). To identify the level of confidence in specific clades reconstructed within the tree, we applied the Bayesian evolutionary analysis by sampling trees method (BEAST2^28,29^) to three randomly selected monophyletic groups of 30 taxa (**Figure 4F**). Broad relationships between the three groups are confidently assigned, while some relationships between very similar taxa are ambiguous, likely requiring additional memory for higher resolution (**Figure 4F**).

Cells exhibiting medium and high edit rate cells clustered together into clades, a feature that is not observed in a scrambled barcode control (**Figure 4E-H**). Scrambling barcodes further significantly impacts the structure of the tree, indicating that tree structure is a result of the correlated barcode patterns generated during the experiment by replicating cells. The mean depth of the reconstructed tree is similar to the number of generations expected for hiPSCs over 23 days of recording, where we would anticipate roughly 23 cell divisions (**Figure 4I**). Together, these results indicate that one-shot transfection has the potential to identify broad clonal features within a population while capturing some detailed lineage relationships across multiple weeks and thousands of cells.

### Hypercascades have the potential to record chromatin transition dynamics

Cellular differentiation and development are accompanied by dynamic chromatin rearrangements, but methods to reconstruct those dynamics in individual cells at scale remain lacking. We reasoned that, if A-to-G base editing were dependent on the chromatin state of its target, it could be possible to make permanent records of chromatin dynamics in conjunction with lineage recording (**Figure 5A**). This is feasible because once edit patterns have been used to reconstruct a lineage tree, it is possible to probabilistically assign individual edits to different points on the tree. The density of edits on any branch of the tree is then directly related to the chromatin state of the target during the corresponding time interval or cell cycle.

**Figure 5:**
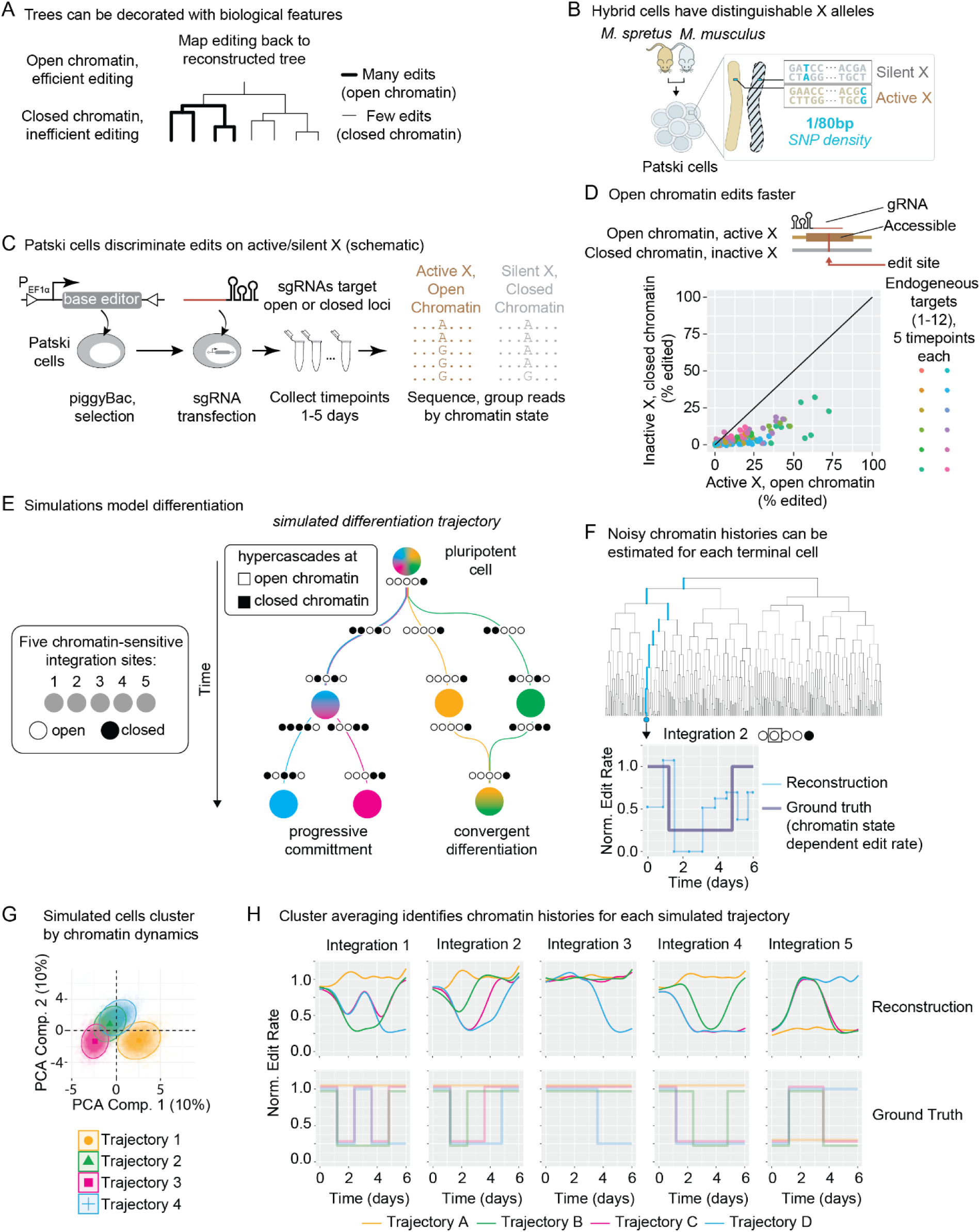
ABE editing is modulated by chromatin state, which has the potential to allow dynamic chromatin recording. **(A)** Linking biological signals, such as chromatin state, to edit rate can enable dynamic inference of the signal in single cells. **(B)** Patski cells, a female hybrid mouse cell line, contain a diploid genome with one allele representing each parental species, rendering alleles distinguishable through single nucleotide polymorphisms. **(C)** One X allele is constitutively silenced, allowing internally controlled investigation of editing at silent and active DNA sequences. **(D)** Across multiple target sites, we estimate a 4-fold increase in editing at open chromatin compared to closed. **(E)** We simulated a hypothetical heterogeneous differentiation, where cells transition through different states (marked by differences in chromatin state across 5 hypercascade-containing loci). **(F)** Lineage reconstruction enables noisy inference of dynamic edit rate modulation (a proxy for chromatin state) over time for each cell. **(G)** Single cells cluster into discrete groups based on their inferred chromatin transition history. **(H)** Averaging across all cells in a cluster gives qualitatively accurate estimates of the dynamic chromatin history experienced by each group.

Cas9^30–32^ and prime^33,34^ editing are dependent on chromatin context. To quantify the effect of chromatin state on ABE activity, we employed a female mouse embryonic kidney cell line derived from two mouse species, M. musculus and M. spretus (the Patski line, **Figure 5B**)^27,35,36^. Patski cells present an ideal system for assaying the effects of chromatin silencing on base editing rates, since identical sequences present on the active and inactive X chromosomes can be simultaneously targeted by a single ABE-gRNA pair but distinguished by a high density of single nucleotide polymorphisms (SNPs) (∼1 per 80 80 bp) (**Figure 5B**). The silent X in this line is associated exclusively with the M. musculus allele, while the active X derives from M. spretus. A recent study of the Patski line characterized differences in chromatin accessibility along the active and silent X chromosomes by ATAC-seq^35^.

We anticipated that chromatin accessibility would differ most strongly near the transcription start site (TSS) of genes where ATAC peaks are present on the active X but not the silent X. If chromatin context affects edit rate, then we would expect to see disproportionate editing at these sites on reads associated with the active X chromosome. To test this, we engineered a stable, polyclonal population of Patski cells constitutively expressing ABE. We designed sgRNAs targeting 12 sites that contained differential ATAC peaks in a 2 kb window upstream of the TSS in 9 separate genomic loci, transiently transfected, and performed amplicon sequencing at multiple time points (**Figure 5C, Supplemental Figure 7A**). We grouped the reads based on their chromatin context. Finally, we quantified the edited fraction at each target site.

We observed pervasive differences in editing between alleles. Almost all target sites exhibited differential editing of the same loci between active and silent X alleles (**Figure 5D**). We identified an average difference of 3.8 ± 0.4 across all genes (**Methods**). Similar results were obtained targeting 10 sites within a single locus (**Supplemental Figure 7B**). These results indicate that chromatin accessibility correlates with edit rate.

Based on these results, we asked whether hypercascade recording could be used to reveal chromatin transitions. We modeled a differentiation process in which a multipotent cell contained 20 distinguishable copies of the hypercascade array integrated at different genomic locations. We simulated a process of progressive fate commitment (**Figure 5E**). 15 integrated hypercascades were assumed to occur in statically open chromatin, which never changes (**Figure 5E**). The remaining 5 were assumed to be dynamic, changing state at defined points along specific lineage trajectories. Open and closed chromatin states were assumed to differ by a 4-fold change in editing rate, similar to that observed in the Patski experiments (**Figure 5D**). The simulation was divided into 5 epochs. Transitioning between each epoch led to progressively more restricted mixing between trajectories, until each differentiation trajectory became canalized into its own path. Two of the trajectories (**Figure 5E**, blue and pink) shared a common progenitor across the first three epochs before diverging to become distinguishable terminal fates. In contrast, the other two trajectories (**Figure 5E**, orange and green) had distinct trajectories from the outset, but converged on the same final fate, modeling convergent differentiation known to occur in some natural systems.

To reconstruct chromatin histories based on the simulated hypercascade edit patterns in these cells, we first reconstructed lineage histories across 200 simulated trees using the UPGMA method. We then estimated the edit rate along every branch of the tree based only on the terminal barcode sequences and inferred lineage using a custom R script. This yielded estimated edit rates over time for each cell (**Figure 5F**). Finally, we averaged over cells with similar reconstructed dynamics to obtain the average chromatin states along each trajectory. To do this, we fit polynomials to the temporal behavior of each integration in each simulated cell, and clustered cells based on their fitting parameters. We used 4th degree polynomials so that multiple transitions between open and closed states could be captured, although in general this could be extended to an arbitrary degree or a Fourier series could be used as appropriate. This method accurately groups simulated cells by trajectory for sufficiently high edit rates (**Figure 5G**). Finally, we used generalized additive model regression^37^ to fit smoothed curves across the cells belonging to each cluster to identify the edit rate across time for each integration locus, mirroring the chromatin accessibility at that locus (**Figure 5H**). All four trajectories yielded curves that reflect ground truth differentiation processes. Together, these results suggest that hypercascades have the potential to be used not only to reconstruct lineage but also to recover chromatin dynamics.

## Discussion

Recording lineage relationships over multiple generations across many cells is necessary to understand organismal development. However, existing recording systems have been limited by small numbers of mutable sites and exponentially decaying edit rates. The hypercascade addresses both challenges by densely packing editable targets in short synthetic DNA sequences, resulting in the generation of, on average, 1 new target site for every edit. (**Figure 1**). In simulations, this strategy dramatically improved reconstruction of lineage relationships in an edit-rate dependent manner (**Figure 2**). Of the many potential hypercascade sequences, we tested three, all of which produced the designed behavior (**Figure 3**), despite occasional out-of-order edits (**Supplemental Figure 1-4**). The hypercascade system is also convenient to use, as all components can be added to cells in a single step (**Figure 4**). Using genomic records from a 23 day recording experiment, we identified edit rate as a clonally inherited feature, and generated a partially resolved tree containing thousands of cells from only a single barcode. In the future, simulations indicate that genomically multiplexing hypercascade barcodes could enable high fidelity reconstruction of developmental processes (**Figure 2**) and chromatin transition dynamics at loci of interest (**Figure 5**).

Engineering the hyperscade requires an accurate editing model. Polyclonal editing experiments (**Figure 3** and **4**) provided estimated rates for out-of-order editing, where the ABE edits targets that have mismatches compared to the gRNA sequence (**Supplemental Figure 1**). Previous studies with wild type Cas9 have shown that the nuclease still edits targets with a small number of mismatches, although at a reduced rate, especially at targets with functional PAM sites.^46,47^ This mismatch editing efficiency has been found to depend on both the sequence of the protospacer and the distance of the mismatches from the PAM site, with higher editing efficiencies for mismatches far from the PAM.^46,47^ As anticipated, edit rates were highest when gRNAs had perfect homology with their target. Also consistent with previous findings, editing rate decreases with more mismatches and is suppressed when the PAM site is broken (**Supplemental Figure 4**). These data allowed us to construct a simplified model of mismatch editing to generate stochastic editing models for each specific hypercascade sequence (**Figure 3D, E** and **4C**, solid lines).

The hypercascade enables new biological recording strategies, including the potential to record chromatin state transition dynamics in parallel across many loci of interest (**Figure 5**). This is only possible due to the high density of targets present in the hypercascade, a key feature of the design that is, to our knowledge, unique to the system: with 70 target sites packed into a single locus, it is feasible to estimate edit rates along each branch of a growing lineage tree at each site of genomic integration (**Figure 5F**). Although we simulated sharp transitions, these were blurred out across time in the resulting estimates (**Figure 5H**). More sophisticated modeling, including incorporation into a Bayesian framework, could enable even better reconstruction of chromatin dynamics. Together, these results suggest that hypercascades have the potential to be used not only to reconstruct lineage but also to recover chromatin dynamics, a possibility that could be explored in future work.

Hypercascade recording has limitations in its current form. At long time scales it is not clear whether barcodes continue to edit effectively. For one of the three targets investigated, full length targets could no longer be recovered by PCR from samples of the population at days 32 and later. Instead, a much shorter product is formed by PCR (**Supplemental Figure 6**). Notably, this target was observed to edit more quickly than the other two targets during the first 15 days, at a rate approximately 5-fold faster. Control samples with the same targets but mismatched gRNAs were still able to be amplified and sequenced; full length targets were observed in the resulting reads (**Figure 3A** and **Supplemental Figure 6**). The other two barcode samples were also able to be recovered and sequenced by PCR amplification (**Supplemental Figure 6**). This could indicate a low rate of barcode collapse due to the repetitive nature of the sequences, which may set an upper bound on the length of time for system recording.

In the future, we anticipate the hypercascade enabling even deeper lineage tree reconstruction with high temporal resolution and extending recording to the level of dynamic chromatin changes. To achieve this, several modifications will be required. First, it will be important to recover edits using single-cell approaches. This would allow us to multiplex the number of hypercascades integrated into each cell and to simultaneously assess individual cellular phenotypes^3^. Second, the number of barcode-distinguishable hypercascades integrated in each cell should be maximized. This is achievable with either multiple rounds of piggyBac transposition and monoclonal selection^3^ or viral engineering approaches^17^. Third, and more technically, further engineering and the use of alternative base editor variants should help to reduce the frequency of hypercascade collapse observed at long time points (**Supplemental Figure 6**). The use of catalytically dead rather than nickase variants of Cas9 in the base editors may help address this issue. Finally, based on simulations, integration of hypercascades at high copy number distributed across the genome should enable capture of chromatin state transitions at each insertion locus (**Figure 5**), potentially yielding a temporal recording of chromatin state that can be recovered with single cell endpoint analysis. Critically, all of these additional steps should be feasible. We therefore anticipate that hypercascades and regenerative editing will open a new epoch of lineage and molecular event recording.

## Acknowledgments

We thank Inna-Marie Strazhnik for expert assistance with preparation of illustrations. We thank Yonatan Stelzer, Amos Tanay, Yoav Mayshar, Netta Reines, Ilana Taieb-Hagerty, Markus Mittnenzweig, Joanna Jachowicz, Magdalenda Zernicka-Goetz, Rong Lu, Giorgia Quadrato, Jean-Paul Urenda, Carla Liacci, Jimmie Ye, Bryson Choy, Mitch Guttman, Matt Thomson, Martin Tran, Donovan Anderson, and members of the Elowitz lab for useful discussions and feedback.

We are grateful to have received support for this project from the Paul G. Allen Frontiers Group and Prime Awarding Agency (award number UWSC10142, M.B.E.), the National Institutes of Health (award numbers R01 MH116508, M.B.E. and training grant T32 GM07616, D.M.C.), the National Science Foundation Graduate Research Fellowship (award number 2139433, D.M.C.), and the Beckman Institute Single-cell Profiling and Engineering Center (SPEC). M.B.E. is a Howard Hughes Medical Institute (HHMI) Investigator. The content is solely the responsibility of the authors and does not necessarily represent the official views of any funding body. This article is subject to HHMI’s Open Access to Publications policy.

## Author contributions

Conceptualization, D.M.C., A.A., A.G., C.L., L.C., M.B.E.; Methodology, D.M.C., R.H., F.H., A.A., A.G., R.B., C.L., L.C., M.B.E.; Software, D.M.C., A.G., R.B.; Formal Analysis, D.M.C., A.G., R.B.; Investigation, D.M.C., R.H., L.K., J.Y.; Writing – Original Draft, D.M.C., M.B.E.; Writing – Review and Editing, D.M.C., R.H., L.K., J.Y., F.H., A.A., A.G., R.B.,C.L., L.C., M.B.E.; Funding Acquisition, D.M.C., C.L., L.C., M.B.E.; Supervision, C.L., L.C., M.B.E.

## Supplemental Figures

**Supplemental Figure 1:**
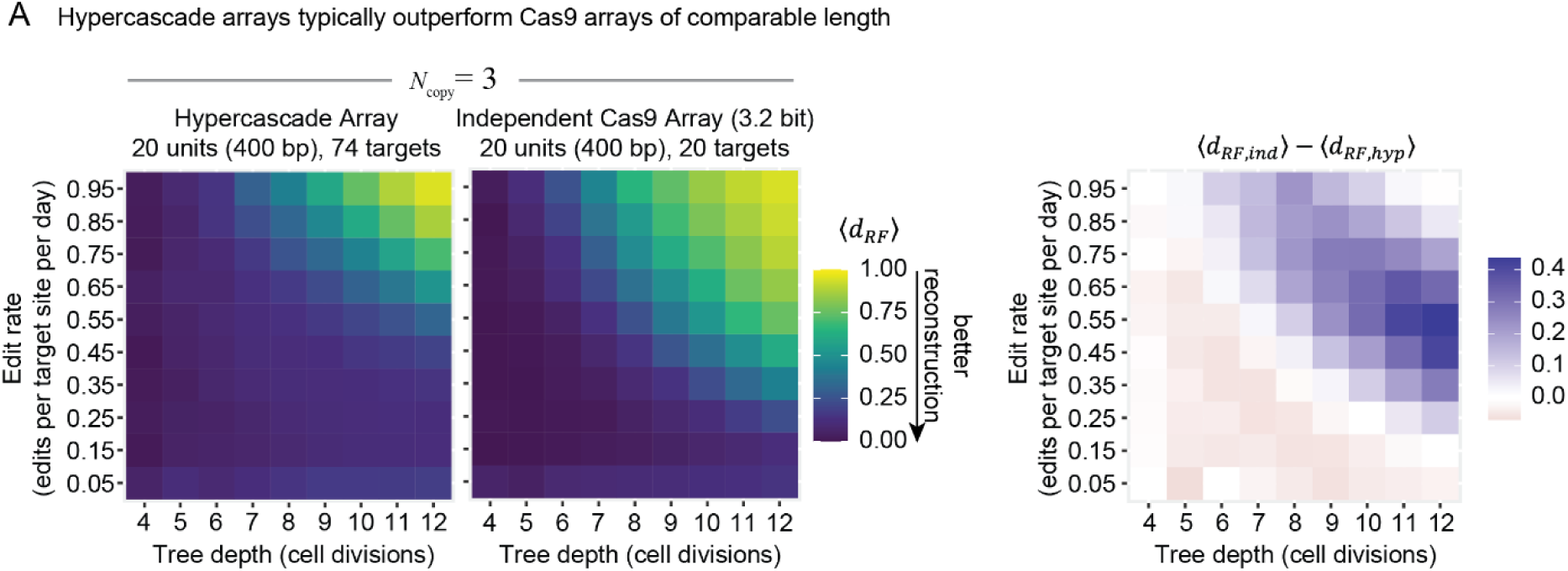
The hypercascade typically outperforms Cas9 based systems of equal length in simulations. Cas9 can produce multiple editing outcomes where base editors produce only a single A-to-G transition. We used a simplified model of Cas9 editing by allowing for each target state to take one of 8 different editing outcomes (see methods). Holding tree depth constant, the hypercascade system outperforms a simulated independent Cas9 target array with comparable sequence length over a variety of edit rates and copy numbers, but underperforms slightly for low edit rates and tree depths.

**Supplemental Figure 2:**
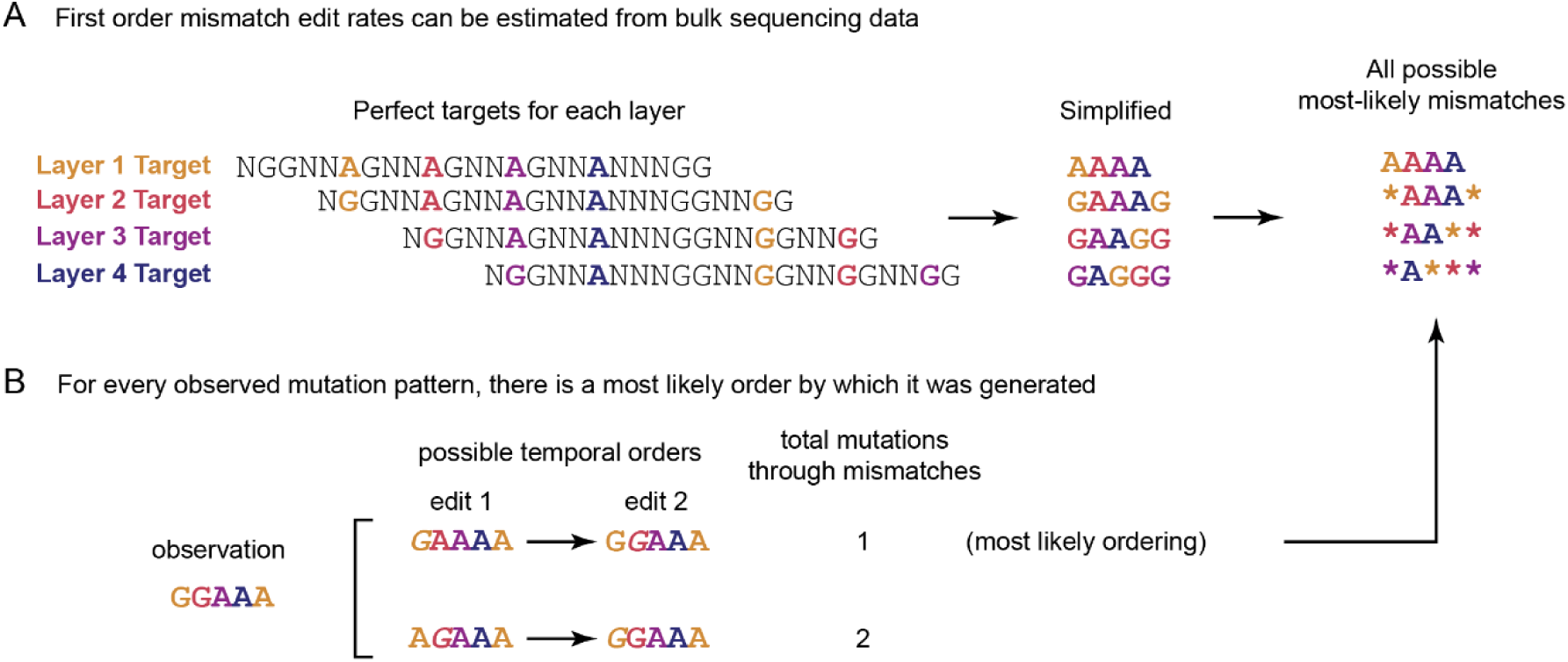
Mismatch edit rates can be estimated. **(A)** Bulk sequencing data can be used to estimate edit rates through either protospacer or PAM site mismatches under several assumptions. **(B)** This is possible by making the assumption that any edit pattern was generated in the most likely ordering of events.

**Supplemental Figure 3:**
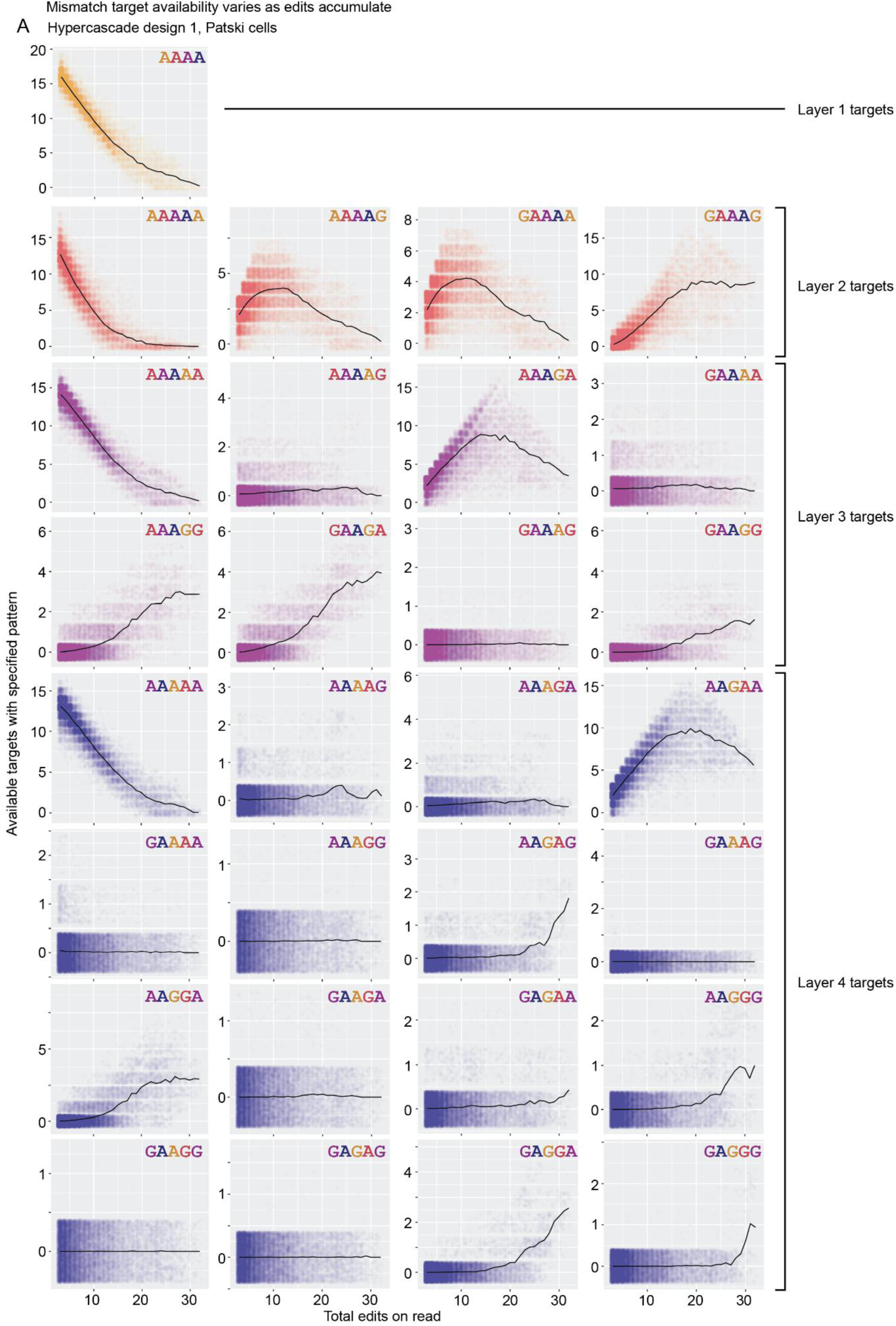
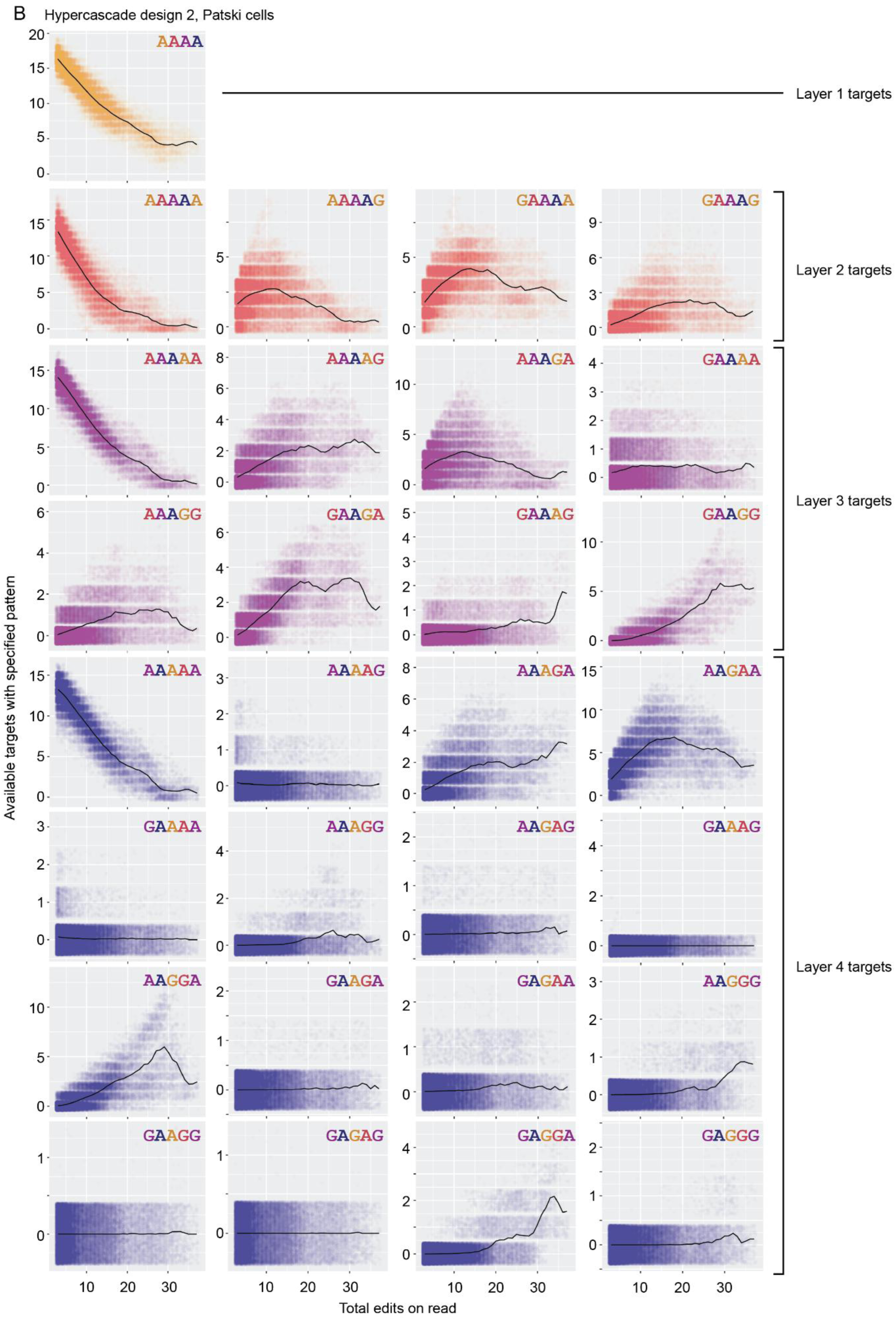
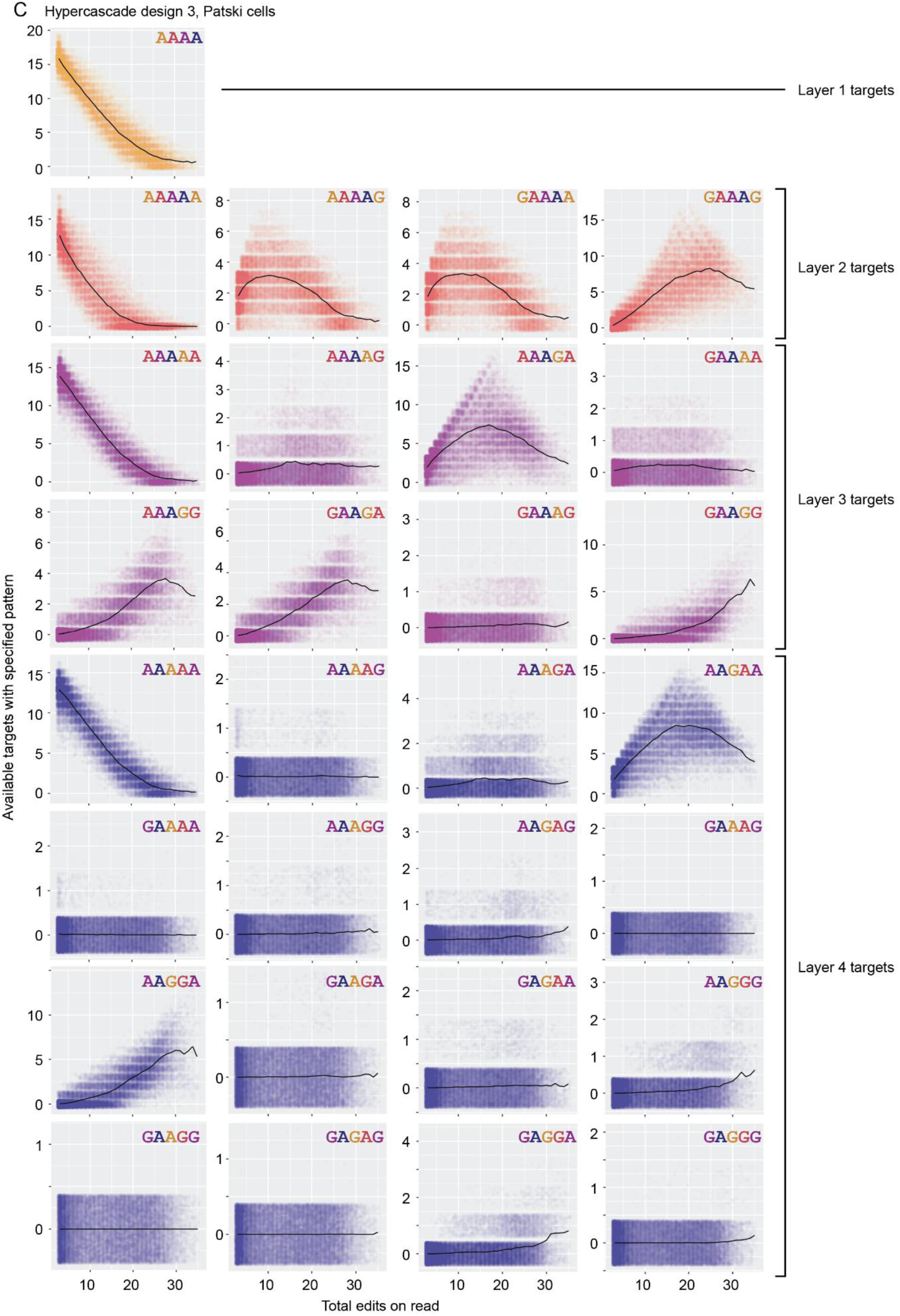
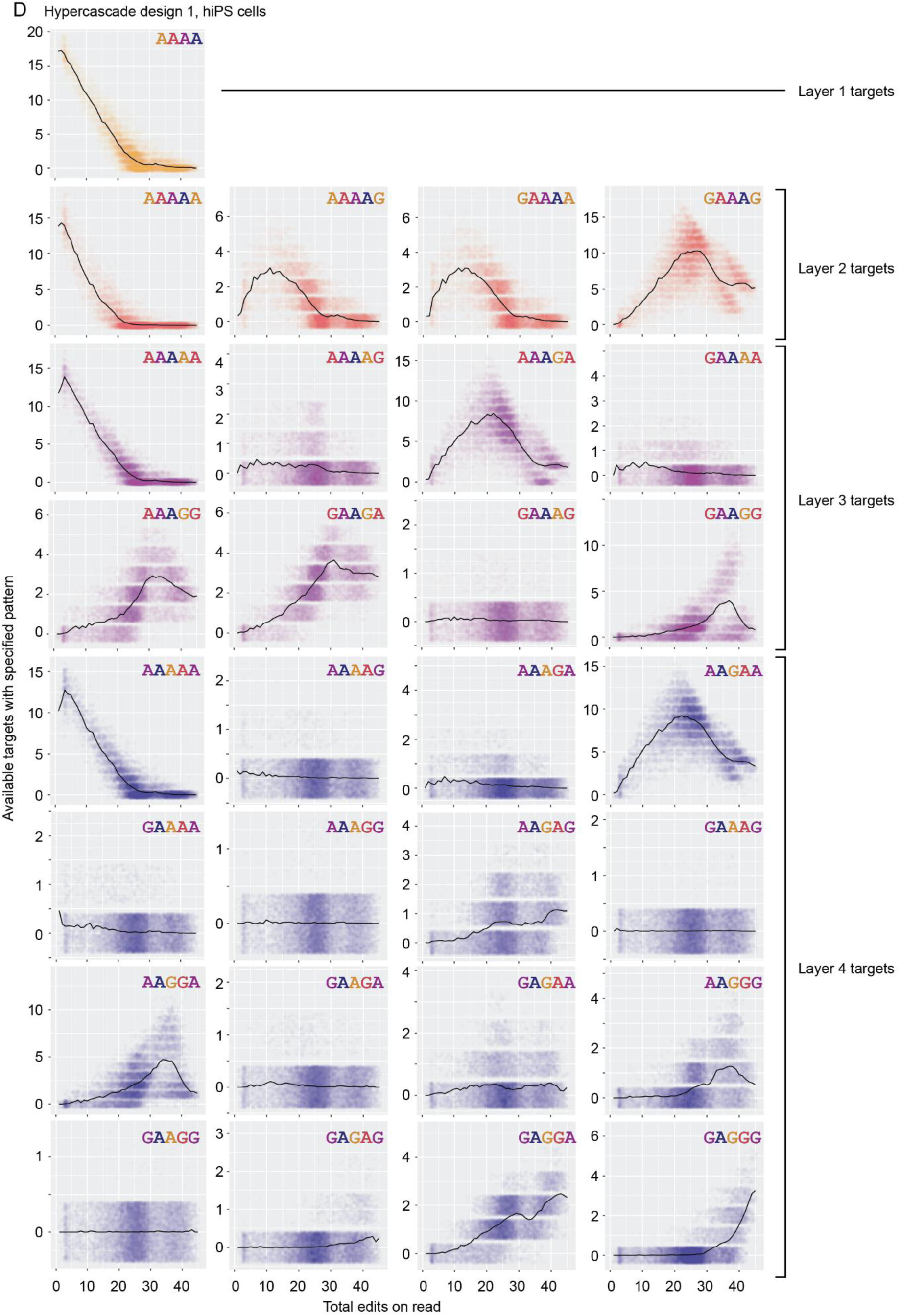
The editing process generates potential mismatch targets dynamically. To estimate mutation rates for a given sequence context, we need to know the number of target sites seen by the editor over time. This can be extracted from bulk sequencing data, both in Patski cells for designs 1 **(A)**, 2 **(B)**, and 3 **(C)**, as well as for design 1 in hiPSCs **(D)**.

**Supplemental Figure 4:**
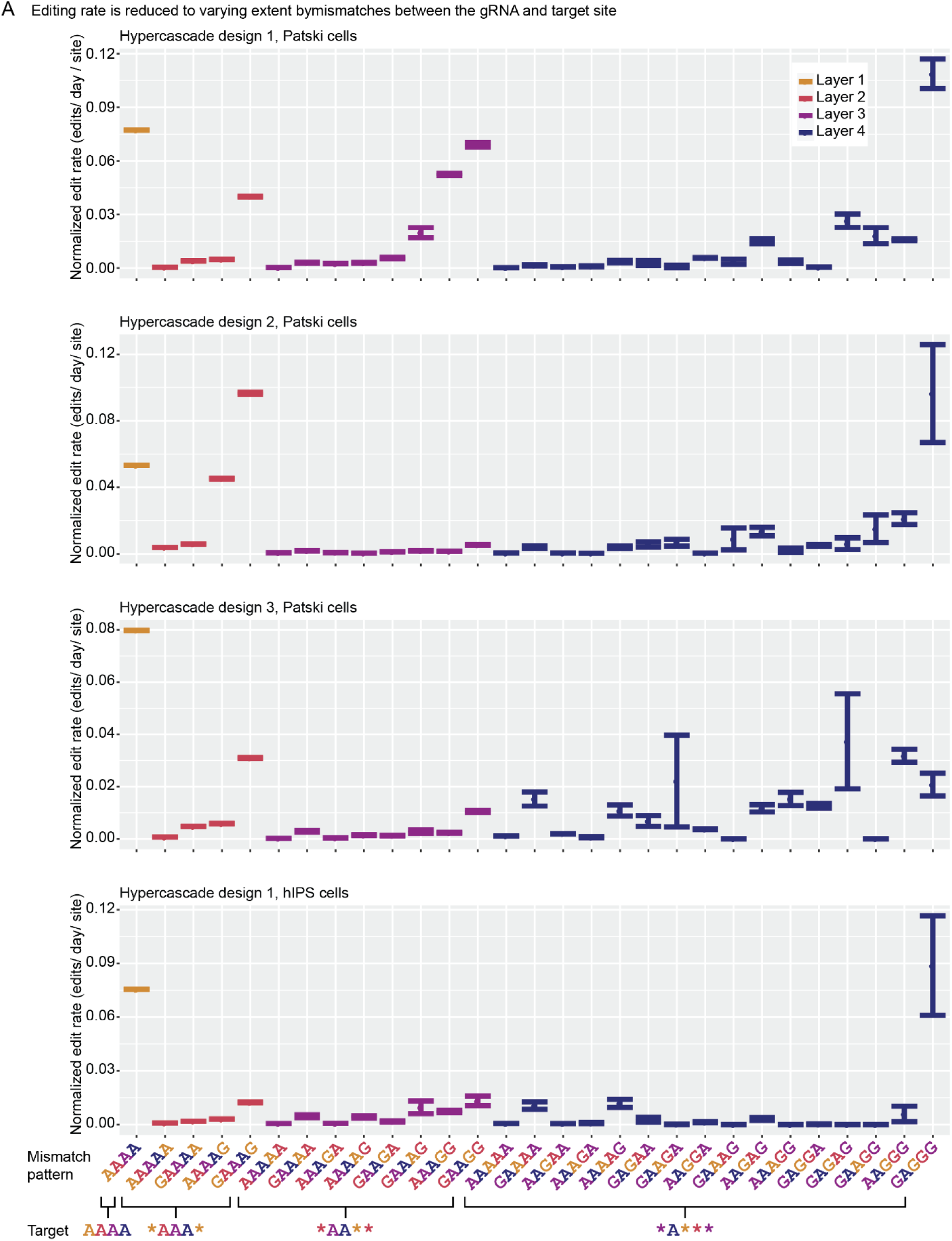
Mismatches decrease the rate of editing across all layers. Mismatch edit rates for many sequence contexts can be estimated from bulk sequencing data. Typically gRNA-protospacer mismatches are estimated to decrease edit rate, with multiple mismatches and mismatches proximal to the PAM site having greater effects. Error bars represent standard error.

**Supplemental Figure 5:**
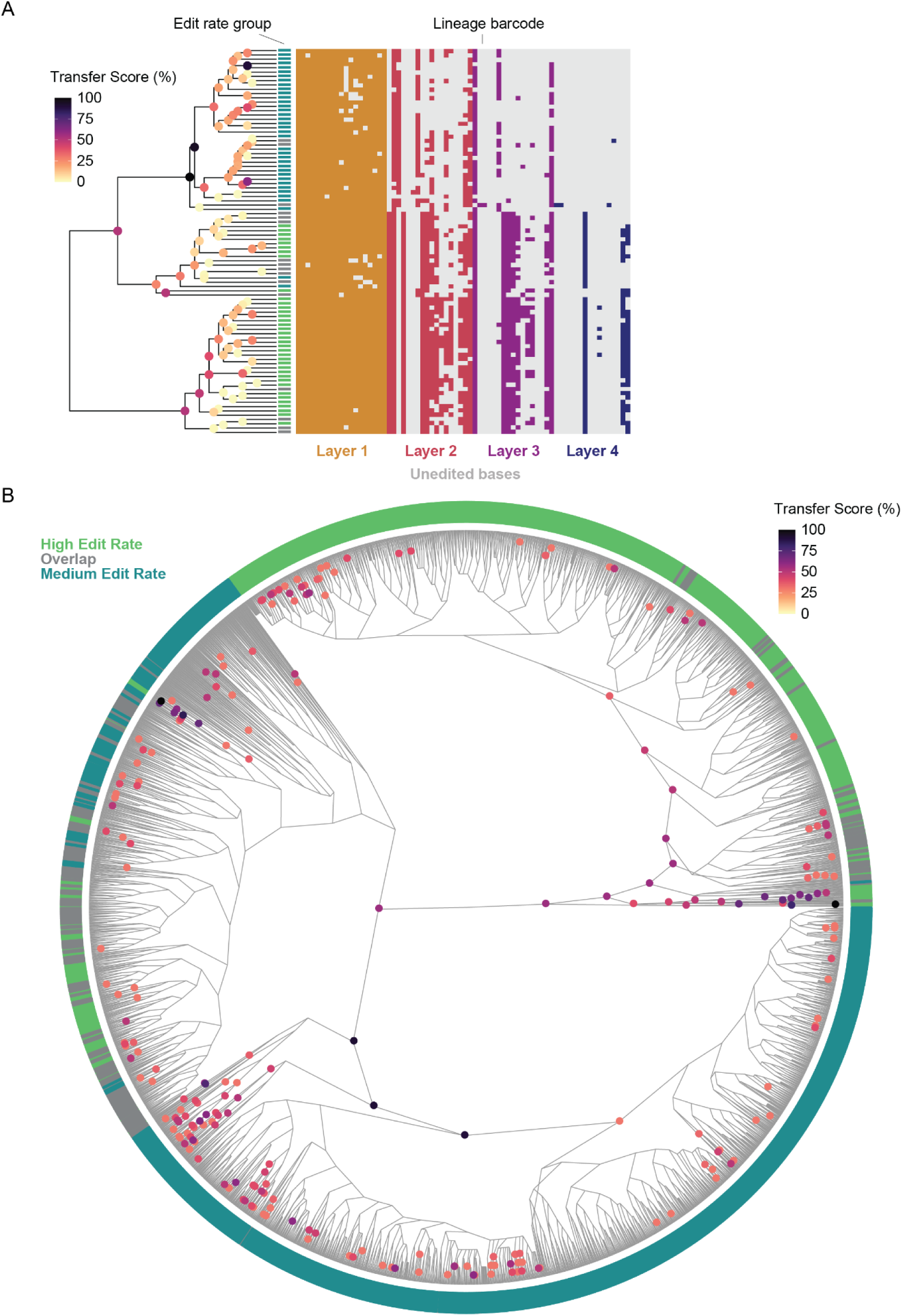
Bayesian analysis and bootstrapping reveal confidently reconstructed phylogenetic relationships among thousands of hiPSCs. **(A)** 90 randomly sampled hypercascades were analyzed using BEAST2 to assess uncertainty in the resulting lineage reconstruction. We identify high transfer scores for a number of clades across the randomly chosen barcodes. **(B)** An alternative method, phylogenetic bootstrapping with UPGMA phylogenetic reconstruction, reveals multiple confidently assigned clades across the full tree. In part **(B)**, only clades with at least 30% bootstrap transfer score are visualized for clarity.

**Supplemental Figure 6:**
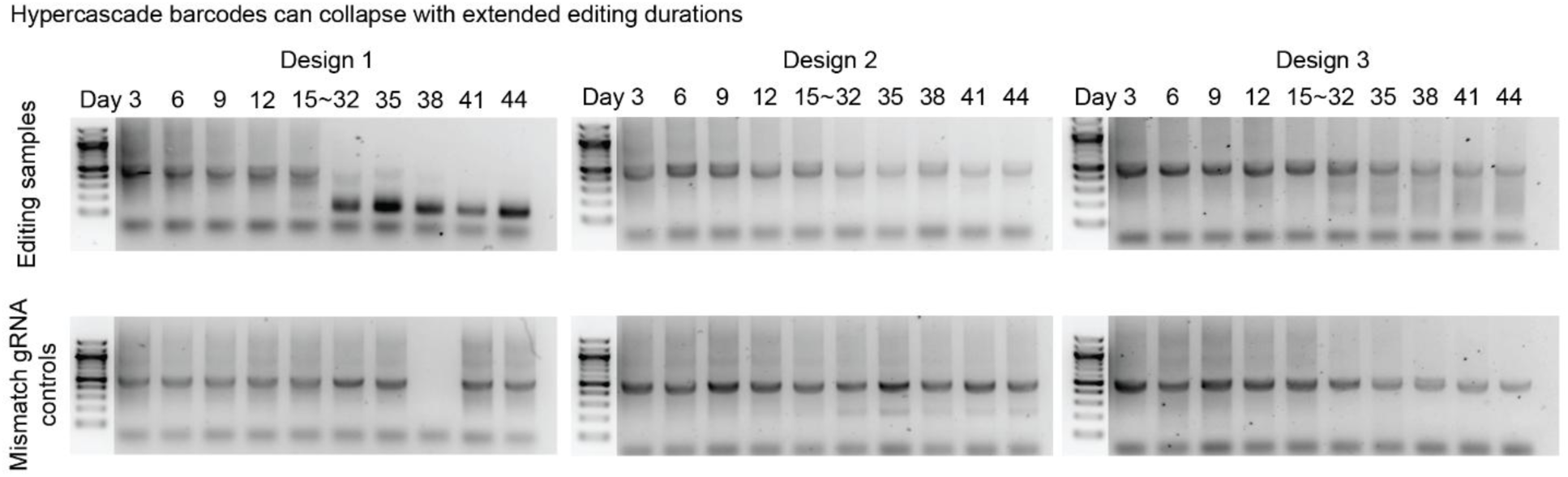
Editing over long times can lead to target array collapse. Hypercascade sequences are amplified from bulk genomic DNA and expected to produce a 500 bp product.

**Supplemental Figure 7:**
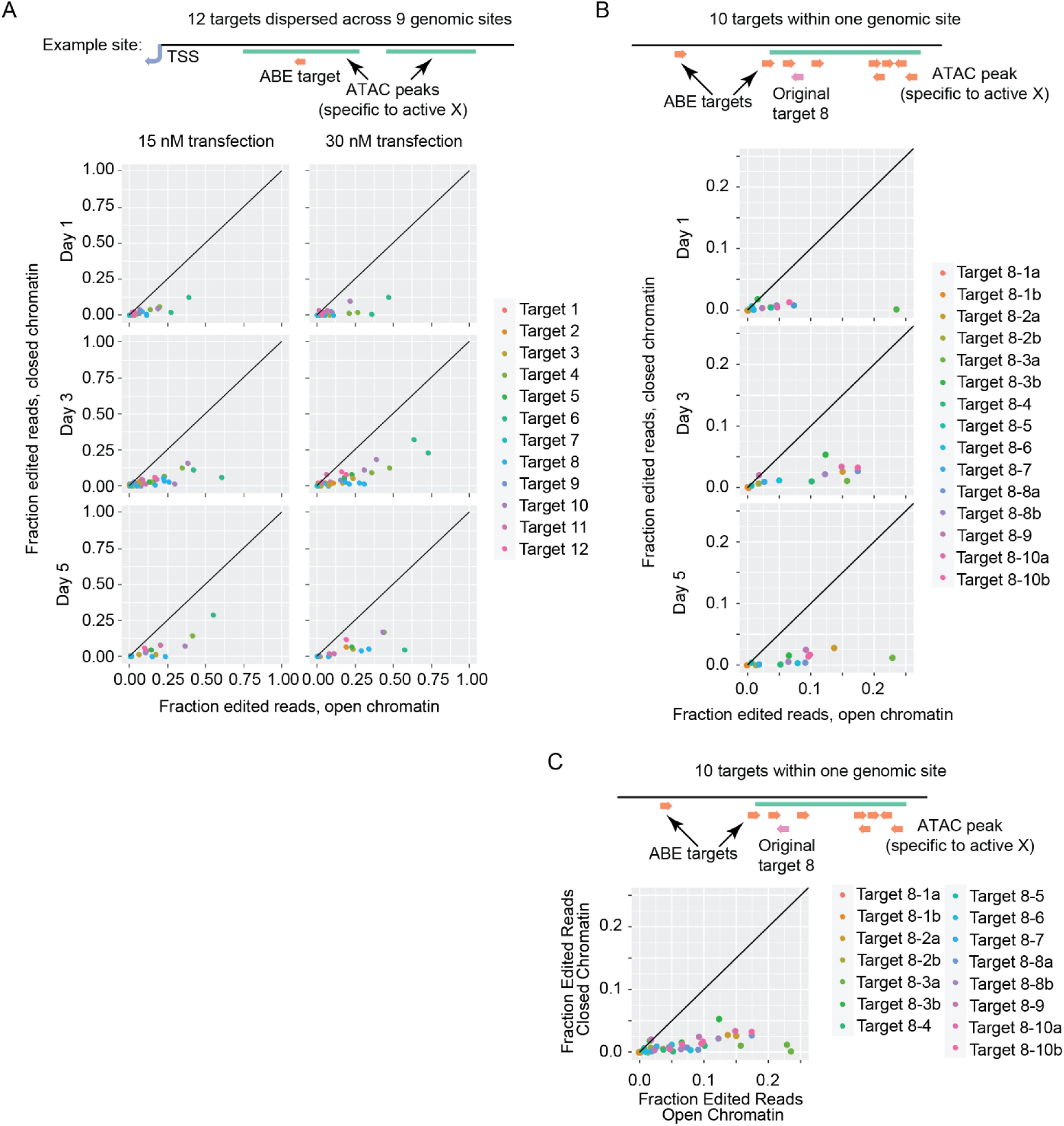
Open chromatin edits more rapidly than closed chromatin. **(A)** 12 targets dispersed across the X chromosome were targeted for base editing over a 1-5 day time course, transfecting either 15 or 30 nM of gRNA into cells constitutively expressing ABE. Targets were selected to have differential chromatin accessibility on the active and inactive X alleles based on existing ATAC sequencing data. Editing was quantified for each allele to determine whether chromatin context impacts ABE edit rate. **(B)** An additional 10 targets were selected and targeted for editing within and around a single differential ATAC peak from panel A. Cells were transfected with 30 nM gRNA only in this case.

**Supplemental Table 1:**
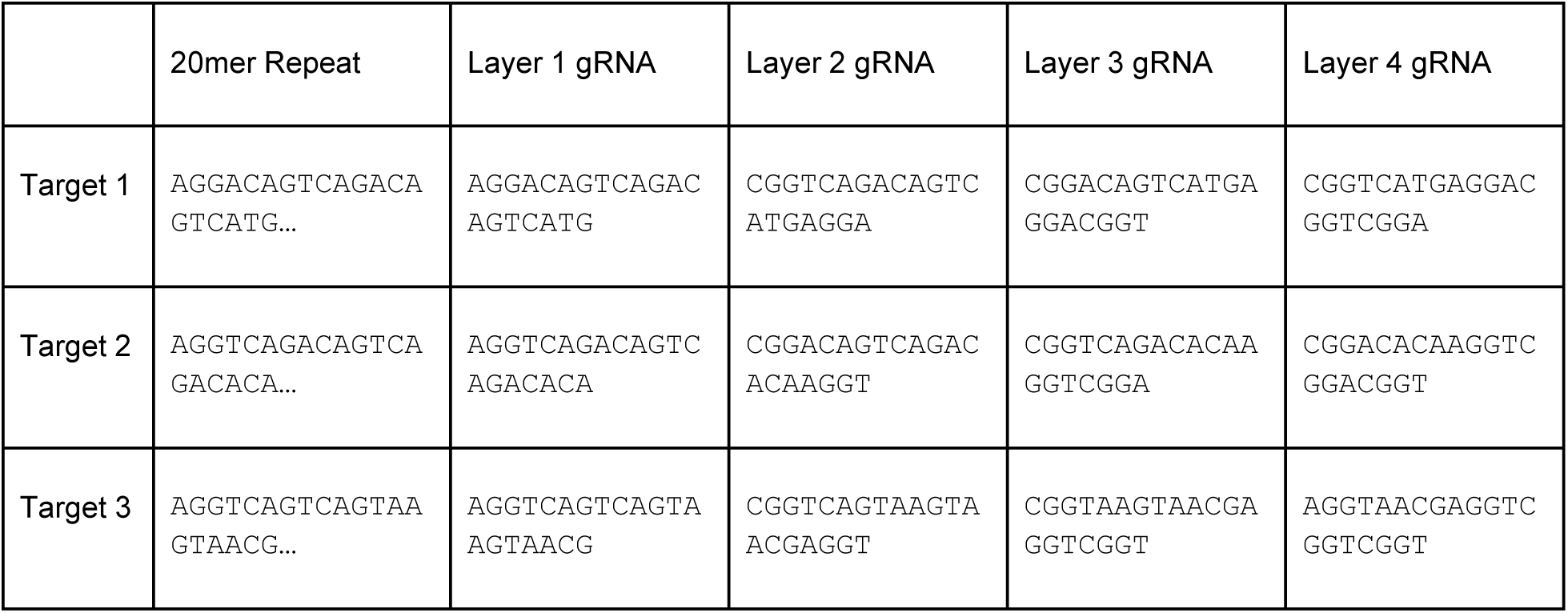
Hypercascade target sequences investigated in this study.

**Supplemental Table 2:**
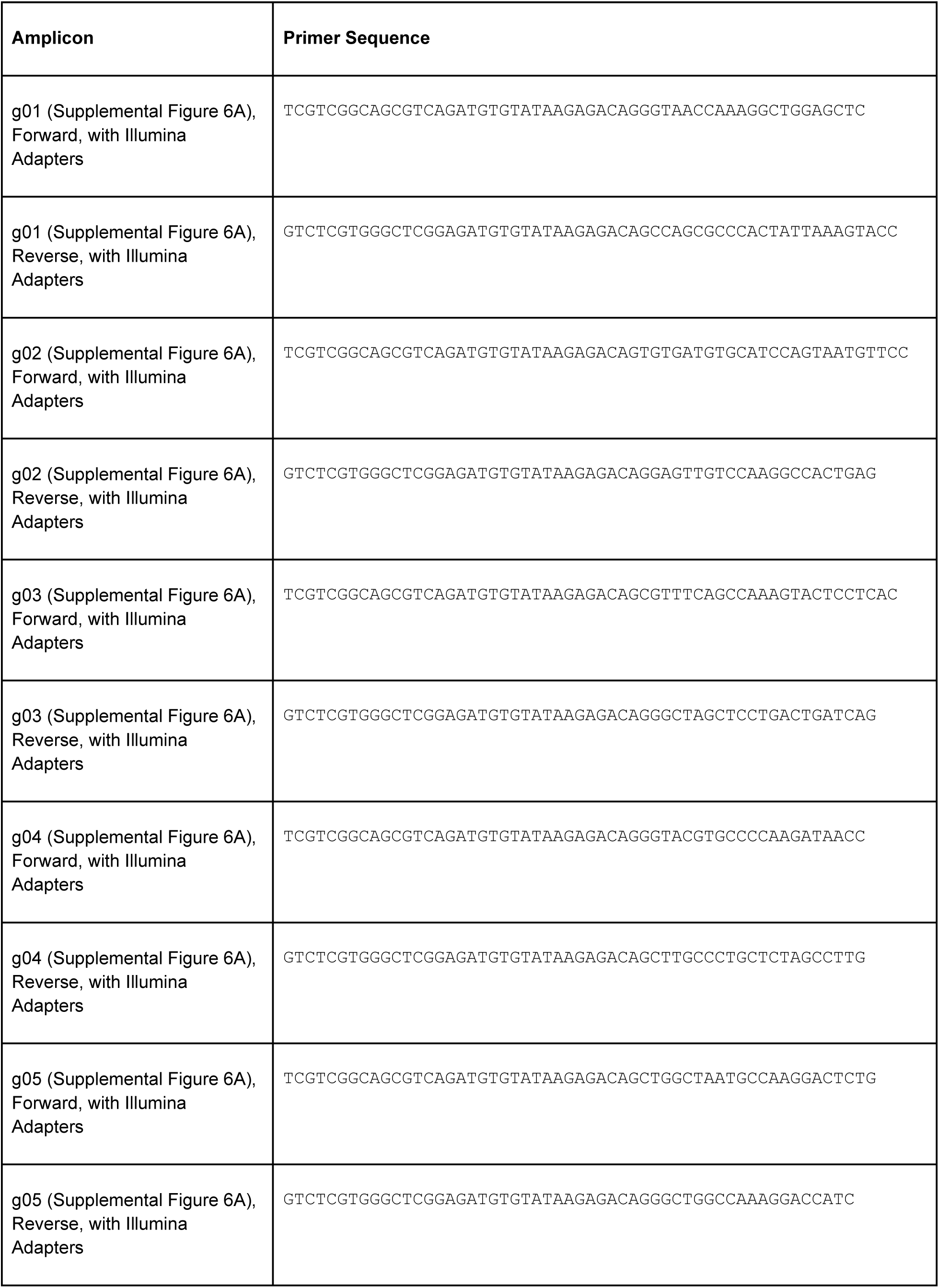
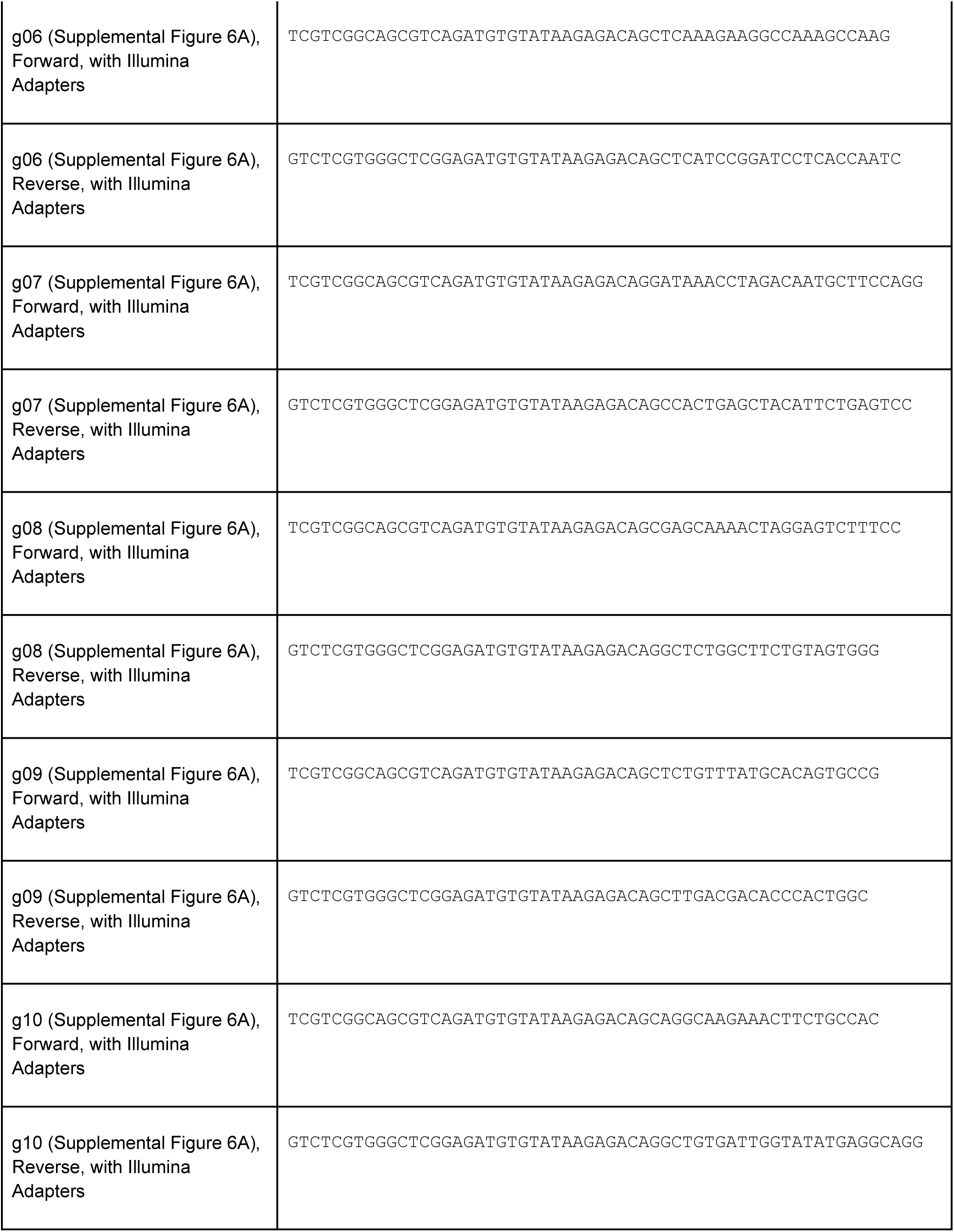
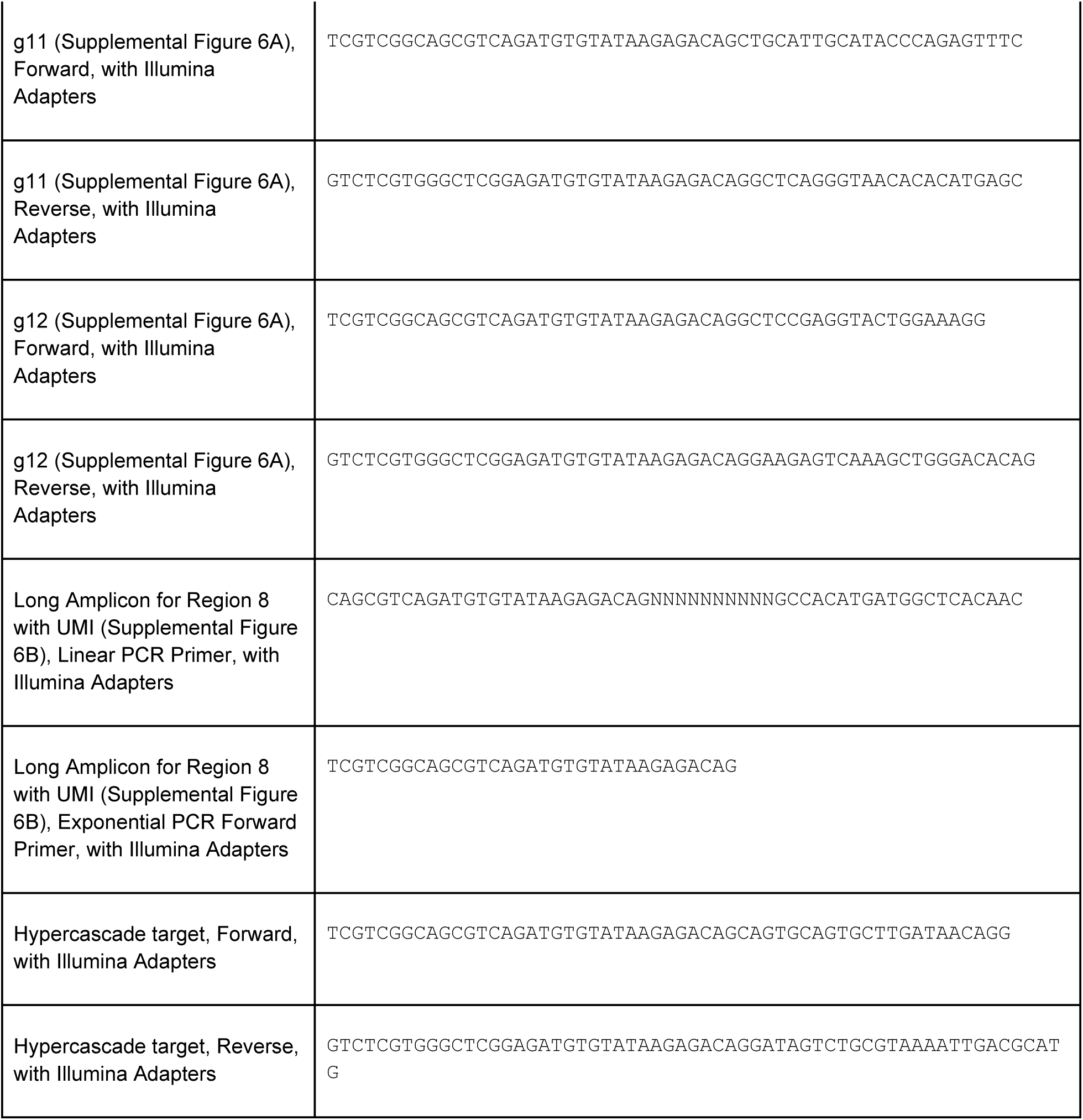
Primer sequences used in this study.

**Supplemental Table 3:**
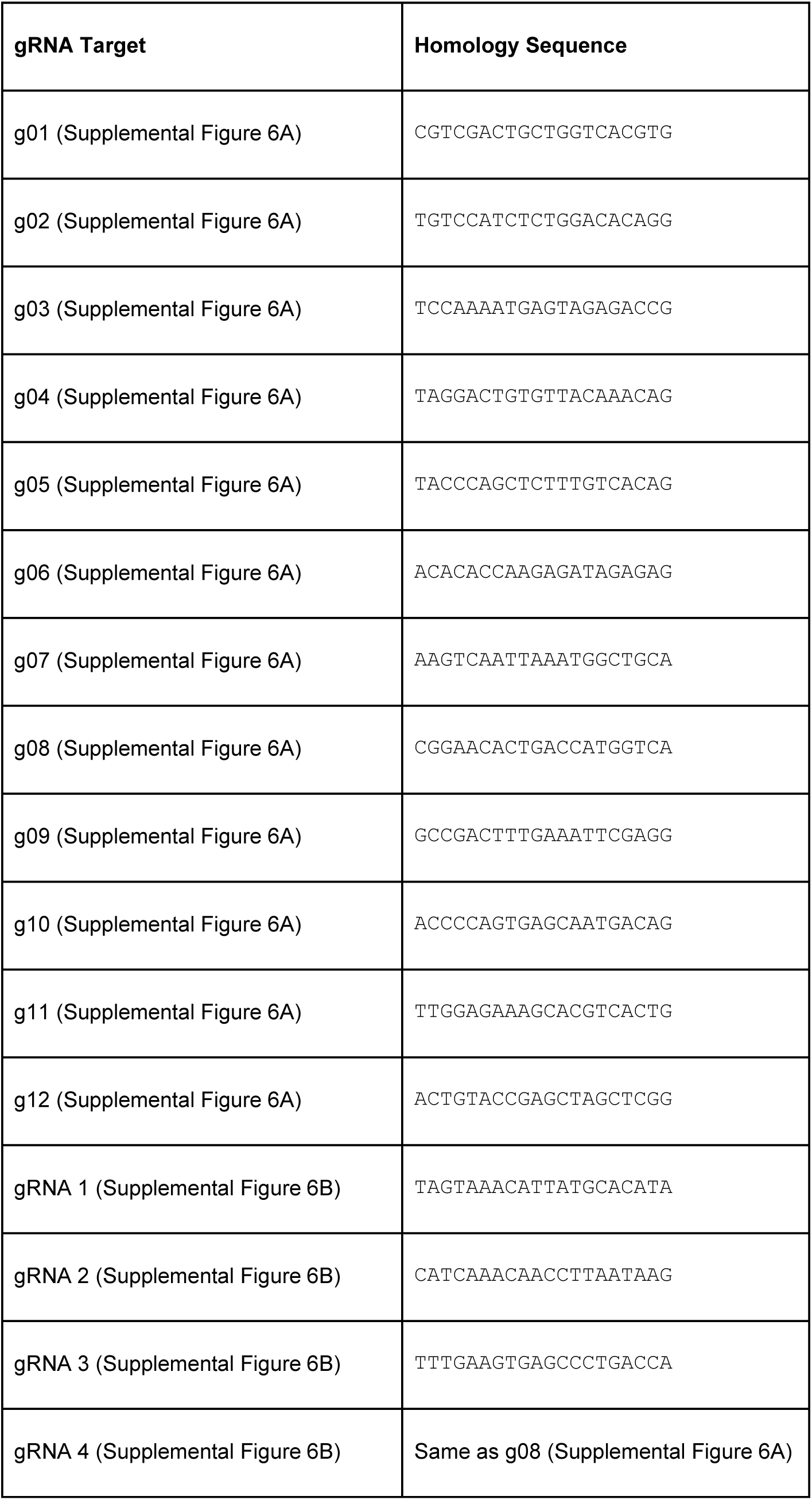
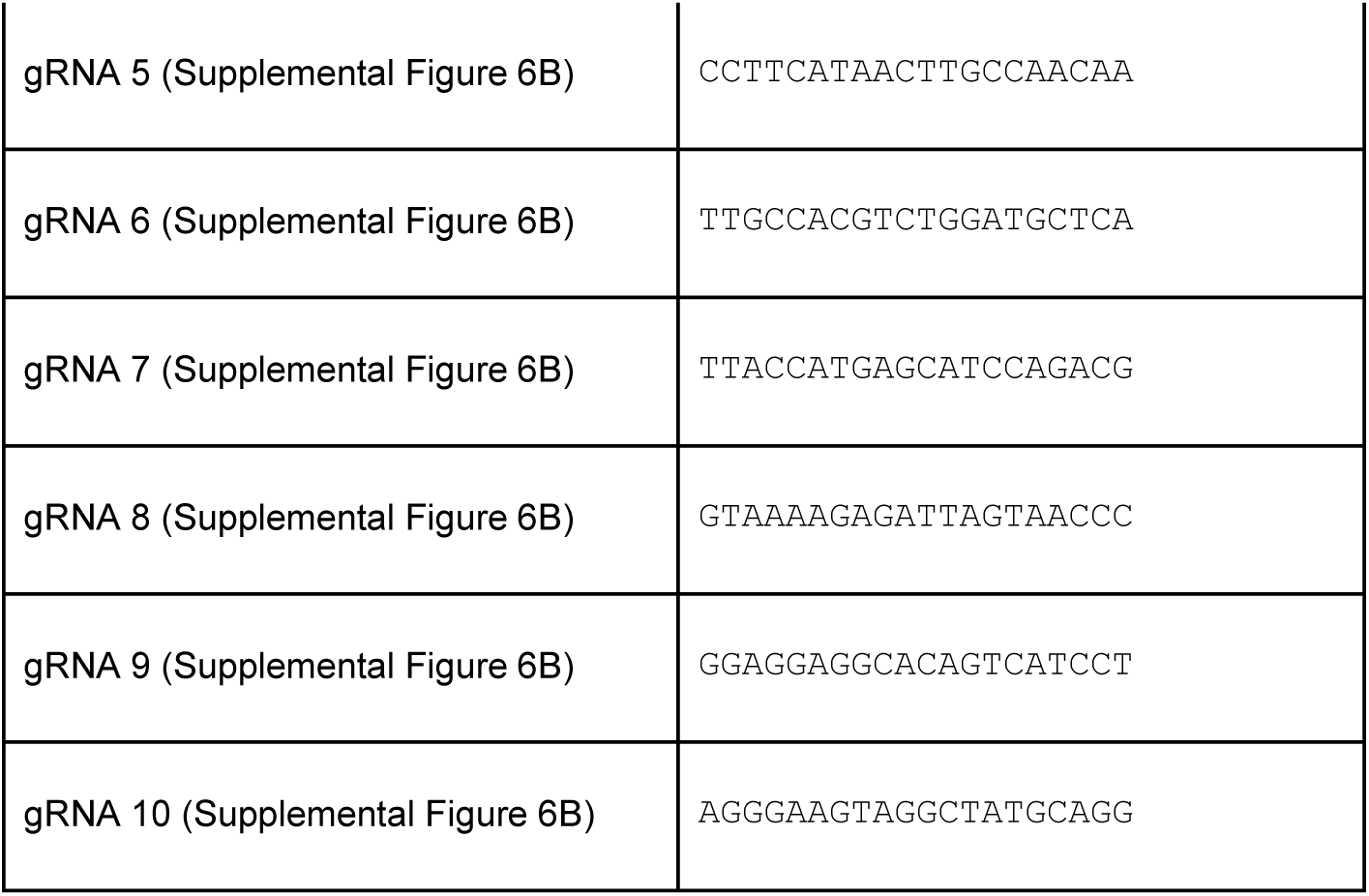
Homology sequences for Patski guide RNA targets (Figure 5D and Supplemental Figure 6).

## Methods

### Cell culture

Patski cells were a generous gift from Chrisitne Disteche and Mitch Guttman. For routine culture, Patski cells were maintained in DMEM with 4.5 g/L glucose, L-glutamine, and sodium pyruvate supplemented with 10% Tet-approved FBS (Clontech) and 500 ug/mL Pen-Strep. When cells reached roughly 80% confluence they were split at a ratio of 1:6 by dissociation with StemPro Accutase (ThermoFisher Scientific). Cells were frozen as necessary in culture medium with 10% DMSO.

Human iPSCs (line WTC-11 / GM25256, NIGMS Human Genetic Cell Repository) were cultured as colonies on embryonic stem cell qualified Matrigel (Corning) with mTeSR1 Complete media (Stem Cell Technologies), changing media daily during culture. Plates were coated according to manufacturer instructions. Cells were passaged every 4-5 days as small clumps using ReLeSR gentle passaging reagent (Stem Cell Technologies).

### Cloning

PiggyBac vectors for ABE integration were developed by Gibson cloning a linear PCR fragment containing ABE7.10 (generated from pCMV-ABE7.10, a gift from David Liu; Addgene plasmid # 102919) into a linearized piggyBac^38^ insertion vector backbone containing the human Ef1a promoter and puromycin resistance (final vector PB-EF1a-ABE-puro).

gRNA expression plasmids were cloned using the GoldenGate^39^ method to simultaneously integrate multiple U6-gRNA fragments in tandem into a piggyBac backbone containing inverted terminal repeats (ITRs) and a blasticidin resistance gene (final vectors H01-4xgRNA-blast, H05-4xgRNA-blast, H10-4xgRNA-blast).

Hypercascade target plasmids, containing repetitive target sites, primer sites, and piggyBac ITRs were synthesized by Genewiz and used directly (final vectors H01, H05, and H10). We elected to use rearranged ITR sequences described previously^40^ to minimize the amount of genomically integrated DNA. In final hypercascade barcode designs, the free 11bp (**Figure 1E, F**) were chosen based on a machine learning algorithm (Azimuth 2.0^26,39,41^) that predicts the editing activity of Cas9 target sites. Each target contained 20 repeats of the 20-mer repeat sequence for a total of 70 editable target sites per barcode (as no PAM site was included for the 20th repeat). Azimuth 2.0 on-target targeting score predictions were chosen to meet the following criteria: layer 1 > 0.7; layer 2 > 0.65; layer 3 > 0.65; layer 4 > 0.6. Final target sequences used in experiments are listed in **Supplemental Table 1**. All plasmids were verified by Sanger sequencing prior to use. Sequence maps are available as supplemental data.

### Generation of stably integrated lines

The Super PiggyBac Transposase system (System Biosciences) was used to generate Patski cells expressing ABE. Patski cells were transfected with PB-EF1-ABE-puro along with a vector encoding Super piggyBac Transposase^38^ at a mass ratio of 4:1 using FuGENE HG transfection reagent following the manufacturer’s protocol in a 6 well plate (Promega). A negative control was performed by transfecting a plasmid encoding GFP without a selection marker. Polyclonal lines were selected with antibiotic (1 ug/mL puromycin) for a period of 14 days, until all cells had died in the negative controls and cells recovered under selection. Lines were expanded in standard media and frozen down for use as needed. Hypercascade targets were integrated into this cell line by transfecting a 10:1 mass ratio mixture of hypercascade target to PB-PGK-ABE-neo vector along with a vector encoding piggyBac transposase using FuGene as described above, then selected with geneticin (500 ug/mL). To start hypercascade editing in Patski cells, a third piggyBac transfection of a vector containing the 4 gRNAs for hypercascade editing and blast resistance was transfected into the polyclonal hypercascade target lines along with piggyBac transposase using the FuGene reagent as above. All combinations of targets and gRNAs were tested individually. These lines were selected with blasticidin (10 ug/mL) for the first 15 days in culture.

For hiPSCs, cells were single suspended 24 hours prior to transfection, counted, and plated in 24 well plates at a concentration of 26.3k cells per cm2. Rock inhibitor Y-27632 (10 uM) was included in the culture media from this point until the end of the antibiotic selection to facilitate cell survival (StemCell Technologies). hiPSCs were transfected with mixtures of 100 ng transposase, 167 ng EF1a-ABE-puro, 167 ng H01, and 167 ng H01-4xgRNA-blast vectors.

LipofectSTEM lipid transfection reagent was included in transfection mixtures to add a final volume of 2 uL to each transfected well of a 24 well plate as recommended by the manufacturer (ThermoFisher). For editing negative control samples, the H01-4xgRNA-blast plasmid was exchanged for 167 ng of ePB-CAG-H2B-Cerulean. As a negative transfection control, parallel samples were transfected with no vector. Transfected cells were allowed to recover for 48 hrs after transfection, then were selected with puromycin (1 ug/mL) for 24 hrs.

In each case, samples were expanded and split with each passage to save cells for sequencing. Collected cells were pelleted at 800xg for 5 min, the supernatant was aspirated, then frozen at - 80 C to await genomic DNA extraction.

### Endogenous editing in Patski cells

To initiate editing of endogenous targets in Patski cells, gRNA was transfected directly into polyclonal cells selected to express PB-EF1a-ABE-puro as described above. Custom crRNA-XT reagents were purchased and used following manufacturer instructions (Integrated DNA Technologies, homology sequences listed in **Supplemental Table 3**). gRNA complexes were transfected by lipofection using the RNAiMAX reagent (ThermoFisher Scientific) at either 15 or 30 nM final gRNA concentration, using a 2:1 ratio of transfection reagent to gRNA in either 6 or 24 well plates. Cells were passaged the day after transfection. Subsequently, samples were split with each passage to save cells for sequencing. Cells for sequencing were centrifuged at 800 RPM for 5 minutes, the supernatant was aspirated, and pellets were frozen at -80 C to await genomic DNA extraction. This sample collection and passaging process was repeated to collect cells at multiple timepoints.

Targets were selected based on a publicly available dataset containing genome-wide ATAC peaks for the Patski cell line focused only on the X chromosomes^35^. The goal was to identify regions that showed differential accessibility between the two X alleles. Since we are interested in target sites that can be informative of cellular state, we decided to focus on promoters. We retrieved 2000 bp length regions upstream of all promoters on the X chromosome and asked whether they contain differential ATAC peaks between the two chromosomes. To classify peaks, we used the ATAC differential score, which ranges from -0.5 (no differential accessibility) to 0.5 (maximum differential accessibility) as defined by Bonora et al^35^. We counted the number of peaks within each promoter region, summed their score and finally selected promoter regions with the highest score. For the X chromosome, the dataset contained in total 1605 ATAC peaks, of which only 204 have 0.5 scores. Therefore, even the promoters with the highest score showed 1 or 2 peaks at most. We finally inspected the selected regions using the UCSC Genome Browser to confirm that the regions did not overlap with transcribed regions, and chose 12 of the resulting ATAC peaks for further analysis.

### Library preparation

Genomic DNA was extracted using the Qiagen DNEasy Blood and Tissue kit, following manufacturer instructions for cultured cells. In some cases, the process was automated using a QiaCube (Qiagen). DNA concentration was measured by Nanodrop. If the concentration was below the limit of detection for the Nanodrop (5 ng/uL) or samples did not exhibit characteristic 260/280nm absorption ratios, samples were further concentrated and purified using the Zymo Clean and Concentrator kit following manufacturer instructions for genomic DNA.

Amplicons were generated for sequencing by a 2- or 3-step PCR approach based on the Illumina 16S amplicon sequencing protocol. Region specific primers were designed for each target site with added Illumina adapter sequences (**Supplementary Table 2**). PCR reactions were set up using Phusion HF 2x Master Mix with gDNA template concentration 5-20 ng/uL and primer concentration 200 nM for the forward and reverse primers. Reactions were denatured for 30 sec at 98 C, then cycled for 35 rounds with 10 sec at 98 C, 30 sec at primer-set specific annealing temperatures chosen using the NEB Tm calculator tool (https://tmcalculator.neb.com/), and 30 sec at the extension temperature of 72 C. Finally, reactions were incubated at the extension temperature for 10 min and stored at 4 C for future processing.

Products were briefly centrifuged then purified with AMPureXP magnetic beads using a 1:1 ratio of sample:beads and following manufacturer instructions (Beckman Coulter). The purified round 1 PCR product served as the template for a second PCR reaction. Primers for the reaction contained Illumina adapter sequences and Nextera XT i5 and i7 combinatorial indices to allow for sample multiplexing. Reactions were cycled 35 times as above with an annealing temperature of 60 C. Samples were briefly centrifuged and purified using AMPure XP beads with a bead-to-sample volume ratio of 1.2:1.

For the samples used to generate **Supplemental Figure 7B**, unique molecular identifiers (UMIs) were incorporated to control for potential PCR bias. In this scheme, an additional linear PCR step was included prior to round 1 PCR using only a region-specific forward primer containing the Illumina adapter sequence and a random sequence of 10 bases. The reaction was cycled between annealing and extension temperatures for 5 rounds, denaturing DNA template only once at the beginning of the reaction to ensure that each extended primer bound to only one DNA target. In this way, target sequences are linearly amplified with one UMI being attributable to each genomic DNA target present in the original sample. The products of this reaction were purified with AMPureXP beads using a bead-to-sample-volume ratio of 1:1 as described above. This reaction was used as the template for the round 1 exponential PCR reaction, priming with a locus-specific reverse primer and a forward primer designed to bind the Illumina adapter overhang of the linearly extended fragments. From this point, samples were processed as described for the non-UMI samples above.

Samples were run out on 2% agarose gels either independently or in mixtures to assess product purity, and concentrations were measured by Nanodrop. Samples were pooled into 4 nM libraries at an equimolar ratio. These libraries were denatured by addition of 0.2 M NaOH, then mixed with similarly denatured PhiX control DNA to increase library diversity (5% PhiX). Samples were run pair end with either 2×250 cycle v2 or 2×300 cycle v3 MiSeq reagent kits following manufacturer instructions. Final libraries were loaded at a concentration of 4 pM.

### Next generation sequencing and data analysis

Hypercascade amplicon sequences were aligned to reference files using the Burrows-Wheeler bwa-mem algorithm^42^. Only reads mapping to the correct target for each line were considered for further analysis. A custom R-script was used to extract base calls and quality scores for each editing target position in the reads. Base calls with quality less than 10 were treated as “N”. The fraction of edited targets over time was computed as the fraction of G base calls divided by the sum of G and A base calls for a time point (**Figure 3C** and **Figure 4B**). For layer distribution analysis and mismatch edit rate estimation, reads containing “N” were excluded (**Figure 3D**, **Figure 4C-E, Supplemental Figures 3** and **4**). Reads were then grouped by total number of edits (G base calls) on the target sites. Groups with sample size less than 30 were excluded.

A Kallisto^43^-based pipeline was applied to analyze data for **Figure 5D** and **Supplemental Figure 7A**. Raw FASTQ files were directly aligned to a manually generated reference file containing, for each amplicon, four sequences: the unedited amplicon sequence from the mm10 genome (UCSC); the unedited sequence from the *Mus spretus* genome (Ensembl); versions of each sequence containing the predicted edit.

For the data presented in **Supplemental Figures 7B** and **7C**, sequencing of relatively long amplicons (∼570 bp) was performed with 2×300 cycle paired end Illumina sequencing in order to obtain information for all edited positions on each individual read. Stringent quality filtering was applied to retain only base reads that had a quality score greater than 20 at the edited positions as described in the following paragraphs.

Raw paired-end FASTQ files were merged using BBMerge using the xloose setting (Department of Energy Joint Genome Institute). Where applicable, random 10-mer UMIs were extracted from reads using UMI-tools^44^. Merged reads were aligned to a reference FASTA file containing only the unedited *Mus musculus* and *Mus spretus* sequences using GraphMap, an aligner built to work well with error-prone reads^45^. The reference file was manually generated by combining short genomic sequences containing the amplicons of interest from both the *spretus* (Ensembl) and mm10 (UCSC) genome assemblies. SAM files were read into R and reads were queried for base calls and quality scores at editing target positions and species SNP locations.

Reads were deduplicated based on their UMI with UMI-tools^44^. Each read contains 5 SNPs that can be used to distinguish the X alleles. Alleles were classified by considering reads that only contained perfect matches to all 5 SNP positions for one species or the other. Editing was assessed at each target site individually, considering only reads that had a quality score greater than 20 at the target site. Samples were discarded if there were fewer than 30 reads remaining after UMI demultiplexing and quality filtering.

For **Figure 4**, lineage relationships were reconstructed between all unique barcode sequences identified from amplicon sequencing at day 23 of the time course experiment. The lineage tree was inferred by computing a pairwise distance matrix between all barcode strings and applying hierarchical clustering in R (the UPGMA method). To assess confidence in the resulting trees, we applied three approaches: 1) selecting 3 random monophyletic clades of 30 cells each, reconstructing lineage relationships using a Bayesian approach (BEAST2^28^), and computing the transfer distance between the posterior samples and the subtree pruned from the UPGMA result (**Figure 4F**); 2) selecting 90 random cells and applying the same Bayesian approach (**Supplemental Figure 5A**); 3) applying phylogenetic bootstrapping^46^ to the full tree with the original UPGMA reconstruction method and computing the transfer distance^47^ across the bootstrap samples for each clade (**Supplemental Figure 5B**).

XML files specifying all BEAST2 modeling information is provided as supplemental data. Briefly, cell division was modeled as a pure birth process (the Yule model) with birth rate estimated. Character mutation was modeled using the ‘irreversible’ package^29^ (source code available at https://github.com/rbouckaert/irreversible). Since targets belong to logical layers that will edit at distinct rates, we allow the rate to vary across sites through the gamma site heterogeneity model, partitioning the allowable rates into 4 categories. We used a strict molecular clock since we do not expect significant rate variation per branch. Priors were chosen to be uninformative.

#### Mismatch edit rate estimation

To estimate mismatch edit rates, we assume, as in previous work^6,21,48,49^, that there is a constant per-target edit rate for each possible gRNA/target combination (referred to here as an “edit pattern”). We assume that the rate of edit accumulation for an edit pattern is given by

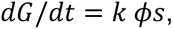

where *G* is the number of edits observed for that edit pattern, *t* is time, *k* is the per-target, per-unit-time editing rate constant (the parameter we are interested in), *ϕ* is a cell line intrinsic parameter reflecting the expression levels of ABE and gRNAs in a cell, and *s* is the number of target sites available for editing. Notably, since we are sampling from a polyclonal population, our measurements are taken from cells with a broad distribution of values. It is assumed that *k* and *ϕ* are independent of time.

Rearranging and integrating, *p* can be estimated by evaluating

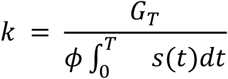

at time *T*, where *G*_*T*_ is the number of edits measured at time *T*. Every read has a *G*^*T*^ associated with it. We also know the total number of edits accumulated on each read and the time when it was measured. We use the overall edit rate per read as an estimate of *ϕ*, normalizing out variance due to heterogeneous ABE and gRNA expression.

It is not possible to assume a functional form for *s*(*t*) a priori due to the interdependent nature of editing. The functional form varies widely for different edit patterns. From the data, we can extract *s* as a function of the total number of edits per read, taking the mean value at each point. Assuming the total number of edits on a read increases linearly with time (an assumption that seems reasonable given the design of the hypercascade and data in **Figures 3** and **4**), we can locate a read along the trajectory of *s* based on the total number of edits on that read, rescale the “edit time” into real time based on the time point when the read was collected, and integrate the mean *s* curve in real time for every read (**Supplemental Figures 2-4**). The mean and standard error of the estimates of *k* over all reads for each edit pattern are plotted in **Supplemental Figure 4**.

### Stochastic simulations

Hypercascade editing was simulated in R using the Gillespie method^23^, where each target site is assigned a propensity to edit in a given time period. Briefly, waiting times between edits are assumed to be exponential. To simulate editing, a waiting time is drawn based on the total propensity to edit any site, then the specific site to be edited is randomly selected, weighting the choice in proportion to each unedited target’s propensity. To investigate theoretical performance of the system, editing was constrained to only allow edits with perfect homology between gRNAs and protospacer sequences as well as completely intact PAM sites (**Figure 2**). For this part of the study, each site was assigned an identical propensity. Cell divisions were modeled by duplicating barcode sequences with waiting times drawn from a distribution derived from Eyring-Stover survival theory that has been shown to better model cell division times than the exponential distribution^50^. We chose to parameterize this distribution using *τ* = 24 and α = 0.9 to model hiPSCs, which typically divide approximately once per day. We screened a variety of overall edit rates, total experiment times, and target array copy numbers, averaging over 30 independent replicates for each parameter set (**Figure 2**).

Independently edited barcodes were simulated in a similar fashion, except without any attenuation of editing propensity based on the states of neighboring targets. To model Cas9-based editing, we augmented the independent model to incorporate 9 different editing outcomes at each target site (all with equal propensities), mirroring the 3.2 bits of information identified on average in Cas9 cut sites^51^.

To better capture editing dynamics in the real system, editing in the presence of mismatches and broken PAM sequences was allowed at an attenuated rate. Further, we allowed each layer its own protospacer-specific edit rate. Mismatch editing was modeled with a simplified, multiplicative model, where each position within the target where mismatches could occur was penalized (**Figure 3D, E**). We applied non-linear least squares fitting to polyclonal editing data to estimate parameters *l*_*k*_ and *p*_*j*_. To fit design 3, we noted lower base qualities than in sequencing other designs, leading to more noise. For this design, we observed a better model fit using only reads with more than 5 total edits accumulated, removing reads that appear to have a small number of edits due to sequencing noise but are actually unedited. For other designs, all reads were used in fitting the model. All simulation and analysis R scripts are available as supplemental data.

### Chromatin state inference simulations

Hypercascade editing was simulated alongside cell divisions as described above, with each cell assigned 20 total barcode arrays, each containing 20 repeating subunits for a total of 74 editable sites per array. 15 of the sites were assigned a constant rate, simulating insertions into statically open chromatin. 5 of the sites had variable edit rate, defined by the chromatin context of the site at different points in time during the simulation (**FIgure 5F**). Editing and cell divisions were allowed to proceed for 6 total days, with open chromatin editing at a rate of 2 edits per barcode array per day and closed chromatin editing 4-fold more slowly (based on empirical measurements from **Figure 5D**).

To estimate the impact of chromatin state on edit rate, we fit a nonlinear least squares model of editing to empirical data collected in Patski cells (**Figure 5D** and **Supplemental Figure 7C**). Edit accumulation for closed and open chromatin was modeled as in prior work^29^ by fitting Equation 1:

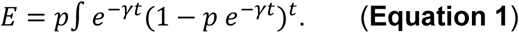

Here, edit accumulation, *E*, is a function of time, *t*, with parameters *p*, the initial probability of editing on transfection of the sample with gRNA, and *γ*, controlling the decay of editing due to degradation and dilution of the gRNAs, which is assumed to be exponential. Parameters were fit to editing time course data (**Figure 5D** and **Supplemental Figure 7C**) to determine the empirical edit accumulation rate for each target using the “nls” function from the “stats” package^52^ in R. The editing ratio between open and closed chromatin was computed by taking the ratio of the estimates of the *p* estimates for *spretus* and *musculus* samples respectively. The standard error for each estimate was propagated to yield the final error term on the editing ratio.

Simulations were divided into 5 evenly-spaced epochs (**Figure 5F**). Four cell-state trajectories were defined over the course of the epochs, with dynamic chromatin trajectories assigned to each state. In some epochs, trajectories had redundant states (for example, in the first epoch all trajectories had the pattern open-open-open-open-closed, as visualized in **Figure 5E, F**). At each cell division, progeny had a chance to switch trajectories based on a predetermined cell state transition matrix, which itself changed in each epoch. Transitions were only allowed between trajectories while they had identical chromatin patterns.

At the end of the 6-day period, lineages were reconstructed based on the resulting barcodes for each cell using UPGMA. Ancestral barcodes were inferred by taking the intersection of edit patterns for each child, starting at the leaves and working back to the root of the tree. Edit rates for each edge were estimated from these ancestral sequences, yielding noisy estimates of edit rate across time for each cell (**Figure 5G**).

To denoise these estimates, edit rate data for each cell was fit to a 4th degree polynomial. Fitting coefficients were clustered using a 4-center Gaussian Mixture Model (Mclust v6.1.1)^53^. Averaged trajectories across each group qualitatively recapitulated the ground truth dynamic chromatin time course (**Figure 5I**).

## Data availability

All next generation sequencing data is available at the NCBI GEO, accession number GSE314635. All analysis scripts, custom code, and additional supplementary files are available at data.caltech.edu, DOI: 10.22002/d15ek-0dx91.

